# The *Vibrio cholerae* MARTX toxin simultaneously induces actin collapse while silencing the inflammatory response to cytoskeletal damage

**DOI:** 10.1101/526616

**Authors:** Patrick J. Woida, Karla J. F. Satchell

## Abstract

Multifunctional autoprocessing repeats-in-toxin (MARTX) toxins are pore-forming toxins that translocate multiple functionally independent effector domains into a target eukaryotic cell. *Vibrio cholerae* colonizes intestinal epithelial cells (IECs) and utilizes a MARTX toxin with three effector domains — an actin cross-linking domain (ACD), a Rho inactivation domain (RID), and an α/β hydrolase domain (ABH) — to suppress innate immunity and enhance colonization. We investigated whether these multiple catalytic enzymes delivered from a single toxin function in a coordinated manner to regulate intestinal innate immunity. Using cultured IECs, we demonstrate that ACD-induced cytoskeletal collapse activated ERK, p38, and JNK mitogen-activated protein kinase (MAPK) signaling to elicit a robust proinflammatory response characterized by production of interleukin-8 (IL-8) and expression of *CXCL8, TNF*, and other proinflammatory genes. However, RID and ABH, which are naturally delivered along with ACD, blocked MAPK activation via Rac1 and thus prevented the ACD-induced inflammation. RID also abolished IL-8 secretion induced by heat-killed bacteria, tumor necrosis factor, and latrunculin A. Thus, MARTX toxins utilize enzymatic multifunctionality to silence the host response to bacterial factors and to the damage it causes. Further, these data show how *V. cholerae* MARTX toxin suppresses intestinal inflammation and contributes to cholera being classically defined as non-inflammatory diarrheal disease.

## INTRODUCTION

Multifunctional autoprocessing repeats-in-toxin (MARTX) toxins incorporate multiple enzymatic functions to promote the virulence of various *Vibrio* species. MARTX toxins are secreted as single 3500 – 5300 amino acid (aa) polypeptides that contain conserved glycine-rich repeats at the N- and C-termini that flank multiple arrayed effector domains and an autoprocessing cysteine protease domain (CPD) (*1*). The glycine-rich repeats are proposed to form a pore in the plasma membrane of eukaryotic cells to translocate the arrayed effectors and the CPD into the target cell (*2–4*). In the cytoplasm, CPD is activated by binding to host inositol hexakisphosphate (InsP_6_) and then auto-cleaves to free the effector domains from the large holotoxin. The individual effectors then can traffic throughout the cell to identify targets and to perform their catalytic functions (*5–7*) (Fig. 1A). Due to this enzymatic multifunctionality, MARTX toxins have been described as bacterial “cluster bombs” that release multiple cytotoxic bomblets into host cells from a single toxin warhead. Although the biochemical function of many of the effector domains is known (*8*), the additive or synergistic benefit of having all these enzymatic functions delivered on a single toxin has yet to be identified (*9*).

**Fig. 1.**
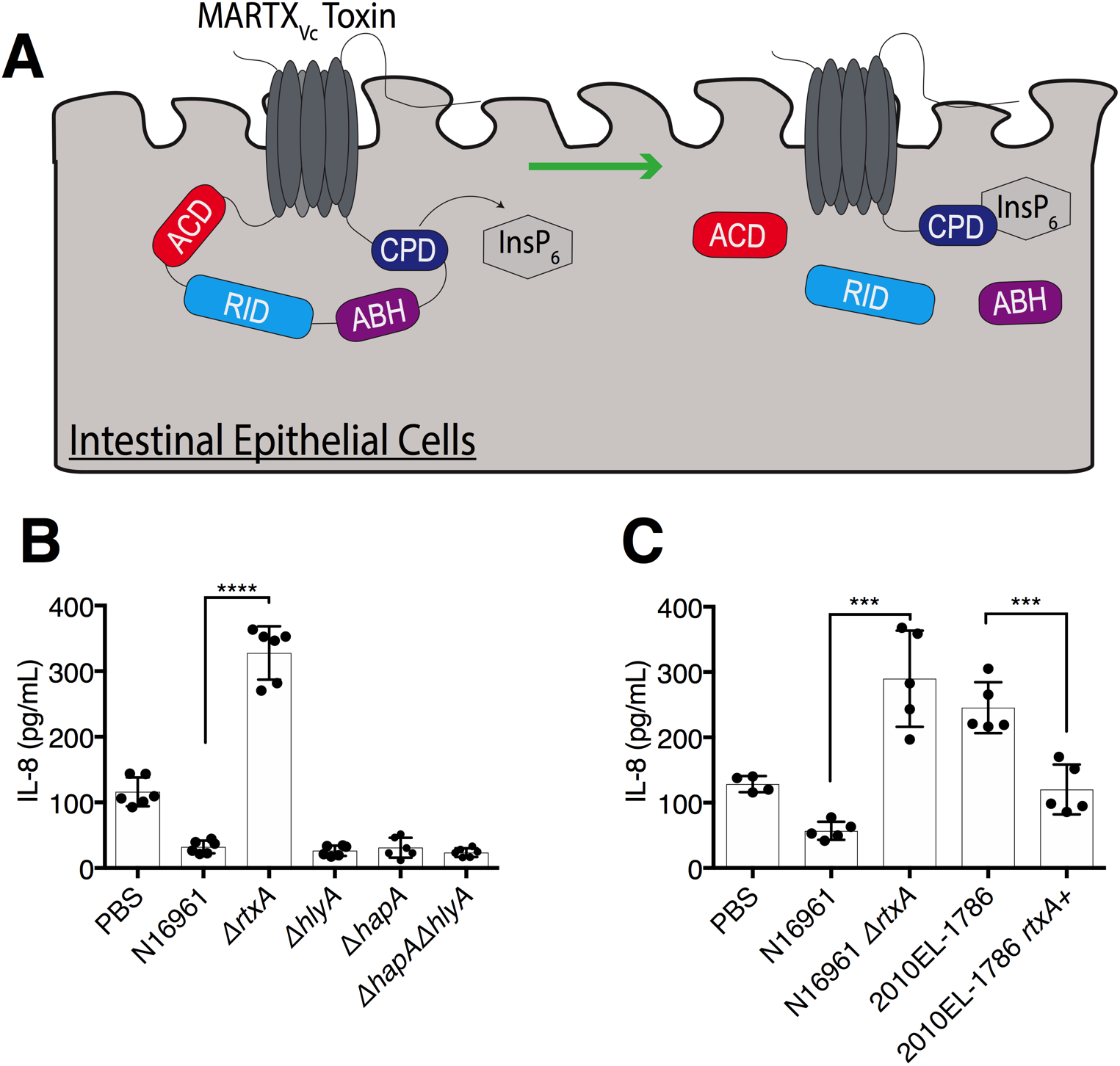
The MARTX_*Vc*_ toxin, and not other *V. cholerae* accessory toxins, suppress IL-8 secretion in IECs. (**A**) Schematic of MARTX*_Vc_* toxin effector domains in IECs. The N- and C-terminal regions of the MARTX*_Vc_* toxin forms a pore in a target eukaryotic cell membrane and translocates the central effector domains (ACD, RID, and ABH) and the protease CPD into the host cell. CPD binds to InsP_6_, which activates the domain’s autoproteolytic activity to separate and release the three effector domains from the holotoxin. (**B, C**) Quantification of IL-8 secretion into the medium by T84 human IECs inoculated with the indicated *V. cholerae* strains (B) or in a clinical isolate with a natural premature stop codon in MARTX*_Vc_* (C). Data for B,C are pooled data from *n* = 3 independent experiments reported as means ± standard deviation (s.d.). Statistical significance of indicated sample pairs determined using Student’s *t*-test (***p<0.001, ****p<0.0001).

*Vibrio cholerae* is the causative agent of the severe diarrheal disease cholera (*10*). In addition to its primary virulence factor, the ADP-ribosylating cholera toxin, pandemic *V. cholerae* El Tor O1 strains secrete a 4,545 aa MARTX*_Vc_* toxin that contributes to enhanced bacterial colonization of the small intestine by protecting the pathogen from neutrophil-mediated clearance during the earliest stages of infection (*11–14*). The early timing of these events suggests that the inhibition of neutrophils and other innate immune cells does not reflect destruction of the cells by the MARTX*_Vc_* toxin, but rather a failure of neutrophils to be recruited to the site of infection. Therefore, the MARTX*_Vc_* toxin might function to limit host signaling that results in innate immune cell recruitment.

Although MARTX toxins can have highly variable effector domain organizations in some species, the MARTX*_Vc_* toxin of nearly all *V. cholerae* strains have the same three effector domains (*15*) (Fig. 1A). The first effector domain is the actin cross-linking domain (ACD) that covalently cross-links monomeric actin to depolymerize actin (*16–18*). The cross-linked actin oligomers also have high binding affinity for formins and other actin-binding proteins, which further disrupts the cytoskeleton (*19*). The second domain is the Rho inactivation domain (RID) that inactivates Rac1 and other Rho-family GTPases by transferring fatty acids onto lysine residues in the C-terminal polybasic region of the GTPases. This acylation of the Rho GTPases blocks them from interacting with downstream effectors (*20*). The action of both ACD and RID results in destruction of the actin cytoskeleton and loss of epithelial cell junction integrity (*2, 20, 21*). The final effector domain is the α/β hydrolase domain (ABH), a highly specific phospholipase A1 that cleaves only phosphatidylinositol 3-phosphate (PI3P), thus inhibiting autophagy and endocytic trafficking (*22*). The activity of ABH also affects the activation state of small GTPase CDC42, possibly due to impact on the phosphorylation state of phosphatidylinositol lipids that control small GTPase activation as PI3P is depleted from the cell (*2, 22*). All three effector domains can also independently disable macrophages, suggesting a mechanism by which the MARTX*_Vc_* toxin protects against clearance of *V. cholerae* from the intestine. However, ACD alone potently inhibits phagocytosis, whereas inhibition by RID and ABH is far less robust (*2*). Therefore, in the context of co-introduction into cells in parallel with ACD, inhibition of phagocytosis is not likely the primary biological function of RID and ABH intoxication. Further, because the MARTX*_Vc_* toxin promotes colonization possibly even well before the onset of inflammation (*11*), inhibition of phagocytic clearance may not be a primary function of the MARTX*_Vc_* toxin.

Intestinal epithelial cells (IECs) act both as a barrier to contain bacteria in the gut lumen and as sensors to detect microbe-associated molecular patterns and release chemokines that recruit immune cells to the site of infection (*23, 24*). Even though *V. cholerae* is typically considered a secretory, non-inflammatory diarrheal disease, these bacteria do elicit mild inflammation in human intestines (*25, 26*). In addition, *V. cholerae* stimulates release of the neutrophil-recruiting chemokine interleukin-8 (IL-8, also known as CXCL8) from human colonic intestinal cells in response to pathogen-associated molecular patterns (PAMPs) including purified flagellin and lipopolysaccharide (*27–30*). However, the extent to which *V. cholerae* induces IL-8 secretion in vitro in response to live bacteria varies dramatically based on the strain isolate (*27, 28*). We hypothesized that the variable response of IECs to live bacteria may reveal that *V. cholerae* actively suppresses proinflammatory signaling induced by PAMPs and that the variation among strain isolates is attributed to strain differences in secreted exotoxins.

In this study, we showe that *V. cholerae* globally suppressed proinflammatory gene expression in IECs and that this signaling inhibition was mediated by the catalytic action of the secreted MARTX*_Vc_* toxin, and not by other accessory toxins. These results explain observed differences in the inflammatory response to various *V. cholerae* strains and why infections with MARTX_Vc_^+^ *V. cholerae* induces a highly secretory, but non-inflammatory, diarrhea. Further, we found that the cytoskeletal destruction initiated by the ACD conversely functioned as a damage-associated molecular pattern (DAMP) that potently induced proinflammatory gene expression through the mitogen-activated protein kinase (MAPK) pathway even more robustly than did *V. cholerae* PAMPs. However, in the context of the MARTX*_Vc_* holotoxin, co-delivery on the same toxin of RID and ABH silenced pathways that would normally transduce the signals to induce proinflammatory gene expression in response to both DAMPs and PAMPs. These data reveal that simultaneous delivery of all three effector domains on a single multifunctional toxin is advantageous as it promotes the ACD-mediated destruction of the actin cytoskeleton without detection of the associated damage by the infected host. Thus, multifunctional toxins can coordinate their multiple enzymatic activities to fine-tune the host response to infection. Further, this study supports that MARTX*_Vc_* toxin suppression of inflammatory gene expression may reduce chemokine and cytokine dependent recruitment of immune cells to the site of infection and may thereby contribute to the differences in inflammation observed between various *V. cholerae* strains (*27, 28*).

## RESULTS

### The MARTX*_Vc_*, and not HlyA or HapA, inhibits IL-8 secretion from IECs

*V. cholerae* secretes three accessory toxins in addition to cholera toxin: the MARTX*_Vc_* toxin encoded by the gene *rtxA*, an α-hemolysin encoded by *hlyA*, and a hemagglutinin/protease encoded by *hapA*. T84 human intestinal epithelial carcinoma cells that have been previously used to characterize the intestinal immune response to *V. cholerae* and other enteric pathogens (*27, 29, 31*). To evaluate whether accessory toxins modulate IL-8 secretion, we treated cultured T84 cells with live *V. cholerae* El Tor O1 strain N16961, which produces all three accessory toxins, or derivative strains with deletions in *rtxA*, *hlyA*, or *hapA* (*32*) at multiplicity of infection (m.o.i.) of 5. Bacteria were removed after 120 minutes (min) and IL-8 was assayed 20 hours later from media containing gentamicin. Whereas N16961 stimulated only small amounts of IL-8 to be secreted into the culture medium, similar to that secreted by untreated control cells (Fig. 1B), N16961Δ*rtxA* induced significantly more IL-8 secretion (Fig. 1B). Bacteria lacking *hlyA* or *hapA*, or both (*ΔhlyAΔhapA*, hereafter referred to as strain KFV119), did not stimulate IL-8 secretion (Fig. 1B). These results reveal that *V. cholerae* has the potential to stimulate release of IL-8 from IECs, but the MARTX*_Vc_* toxin actively suppresses the response.

The atypical El Tor O1 strain 2010EL-1786 from the 2010 Haiti outbreak is known to be more proinflammatory than other stains of *V*. *cholerae* (*33*). This strain has a premature stop codon in *rtxA* that prevents secretion of the MARTX*_Vc_* toxin (*15*). Consistent with results using N16961, 2010EL-1786 induced a significant increase in IL-8 secretion from IECs, but this response was suppressed when the premature stop codon in *rtxA* was restored to a Trp codon (Fig. 1C). These results show that *V. cholerae* activates proinflammatory chemokine secretion that is suppressed in *rtxA^+^ V. cholerae* strains.

### MARTX*_Vc_* toxin effector domains, and not the pore, inhibit IL-8 secretion in IECs

The MARTX_Vc_ toxin has four putative cytotoxic functions: the formation of a pore and the biochemical activities of its three effector domains (*9*). In previous studies (*2*), a derivative of KFV119 was generated in which the *rtxA* gene was modified to replace the sequences that encode the effector domains with an in-frame sequence for β-lactamase (Bla). This strain produces a toxin that forms a functional pore and is able to translocate, but lacks all effector activity (*rtxA::bla*). The individual effector domains ACD (*acd::bla*), RID (*rid::bla*), and ABH (*abh::bla*) were then individually restored into the parent *rtxA::bla* strain to create toxins that delivered one of the three effectors (Fig. 2A-E). Catalytically inactive derivatives of these effectors and a Δ*rtx* negative control strain were also generated (*2*). These strains were fully characterized, including demonstration of similar detection of Bla activity within eukaryotic cells after 60 min incubation of bacteria with cells across all strains (*2*).

**Fig. 2.**
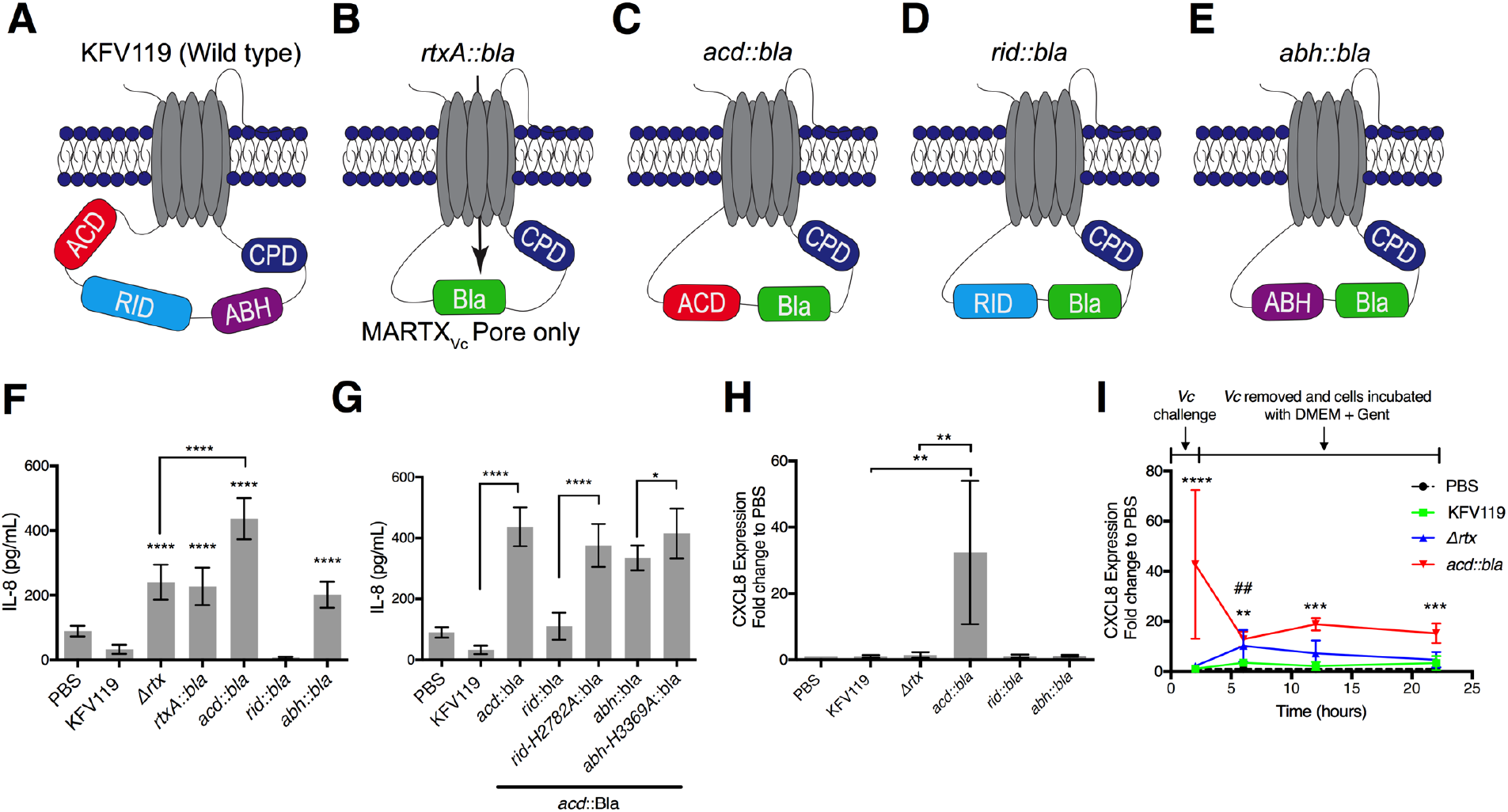
The MARTX*_Vc_* toxin effector domains, and not the pore, differentially regulate IL-8 secretion in IECs. (**A-E**) Schematics of MARTX*_Vc_* toxin effector arrangements in wild-type *V*. *cholera* (**A**) and in strains modified to translocate β-lactamase (Bla) in place of all three effector domains (**B**) and in single effector gain-of-function strains (*2*) in which each effector domain was individually restored (**C–E**). (**F, G**) Quantification of IL-8 secretion from T84 IECs inoculated with one or two *V. cholerae* effector-free Bla and single effector gain-of-function strains inoculated 1:1 (each m.o.i=2.5) as indicated. (**H**) Quantification of *CXCL8* expression by qPCR in T84 cells at 120 min post-inoculation with *V. cholerae* strains as indicated. (**I**) Quantification of expression by qPCR with bacteria removal after 2 hr and then assayed at times indicated from antibiotic-containing media. Data for all panels are pooled from *n* = 3 independent experiments reported as means ± s.d. Statistical significance determined by One-way ANOVA with Tukey’s multiple comparison’s test for samples as indicated (*p<0.5, ** p<0.01, ***p<0.001, ****p<0.0001). For panel I, PBS mock treated sample was normalized to 1 and asterisks indicate multiple comparisons between *acd::bla* and PBS and ## (p<0.001) for multiple comparison between Δ*rtx* and PBS set at the same time point.

The *rtxA::bla* strain induced similar amounts of IL-8 secretion from IECs as did the *Δrtx* strain, but significantly more IL-8 secretion than did KFV119 (Fig. 2F). This result indicates that the MARTX*_Vc_* pore alone does not suppress IL-8 secretion. Unexpectedly, the *acd::bla* strain induced significantly more IL-8 secretion compared to both the *Δrtx* and *rtxA::bla* strains (Fig. 2F). The *rid::bla* strain induced IL-8 secretion equivalent to that induced by KFV119 (Fig. 2F). These results indicate that RID is sufficient to suppress *V. cholerae-*induced IL-8 secretion, whereas ACD actually exacerbates IL-8 secretion beyond stimulation by live bacteria lacking the MARTX effectors (Fig. 2F). In fact, when IECs were simultaneously co-inoculated with both the *acd::bla* and the *rid::bla* strains at equal m.o.i of 2.5 (combined m.o.i of 5), IL-8 secretion was also suppressed (Fig. 2G). This was due to the acylation activity of RID, because the *rid*-H2782A*::bla* catalytically inactive strain did not suppress IL-8. In addition, although less robust, co-inoculation of *acd::bla* with *abh::bla* also attenuated IL-8 secretion (Fig. 2G). These results reveal that RID, and to a lesser extent ABH, inhibits IL-8 secretion induced by ACD.

### The MARTX*_Vc_* toxin inhibits the expression of *CXCL8*

A block to IL-8 secretion can occur by inhibition of IL-8 protein secretion or inhibition of *CXCL8* (IL-8) gene expression. As done above, cells were treated with bacteria for 120 min and gene expression determined from cells collected from antibiotic-containing media. There was no significant change in transcription of the *CXCL8* gene in intestinal cells inoculated with KFV119 or *Δrtx* compared to mock-treated cells (Fig. 2H) at two hours when the bacteria were removed. There was also no significant change due to RID or ABH. By contrast, a significant 32-fold increase was induced by ACD (Fig. 2H). These results indicate that MARTX*_Vc_* effector domain modulation of IL-8 secretion occurred primarily due to blocking of *CXCL8* gene transcription.

The absence of a response to *Δrtx* was unexpected since *V. cholerae* alone induced IL-8 secretion (Fig. 1B and 2F) (*27*). Since chemokine secretion from cells by ELISA requires a long incubation period following the 120 min bacterial challenge, we hypothesized that the IEC response to *V. cholerae* occurs after exposure to bacteria, during the 20 hour incubation in antibiotic-containing media. Therefore, changes in *CXCL8* expression were measured overtime following bacterial challenge. Confirming previous results, no change in *CXCL8* expression was observed by KFV119 at any timepoint measured after bacteria removal (Fig. 2I). However, while *Δrtx* did not induce changes in expression at 120 min post-inoculation, there was a significant 10-fold increase in expression four hours after bacterial removal (six hours post-inoculation) that eventually returned to baseline between 12 and 22 hours (Fig. 2I). As observed previously, *acd::bla* induced a significant 42-fold change in *CXCL8* expression at two hours. While this response diminished overtime, relative abundance of *CXCL8* mRNA remained 10- to 18-fold higher than mock-treated cells (Fig. 2I). These data further show that ACD-mediated *CXCL8* expression is stimulated as much as four hours before *V. cholerae*-induced gene expression. Therefore, in addition to suppressing the host response to effector activity, *V. cholerae* may utilize the MARTX*_Vc_* toxin to disable host inflammatory responses prior to detection of bacterial PAMPs, such as flagellin and lipopolysaccharide (*29, 30*).

### ACD, but not the MARTX*_Vc_* holotoxin, induces proinflammatory gene expression in IECs

To determine the breadth of the proinflammatory immune response stimulated by ACD, we utilized a whole transcriptome RNA-sequencing (RNA-seq) approach. Cells were treated with various bacterial strains for two hours prior to transcriptional profiling. Statistically significant results were given a −1 ≤ 0 ≤ 1 log_2_ fold change cut off to adjust for biologically significant changes (Fig. 3A and B, Table S1). A single gene *JUN* was stimulated by KFV119 compared to uninoculated cells, but this stimulation was not consistent when validated by qPCR (Fig. 3C). Thus, there was no significant activation of any host gene after two hours treatment of IECs with MARTX*_Vc_*+ *V. cholerae*.

**Fig. 3.**
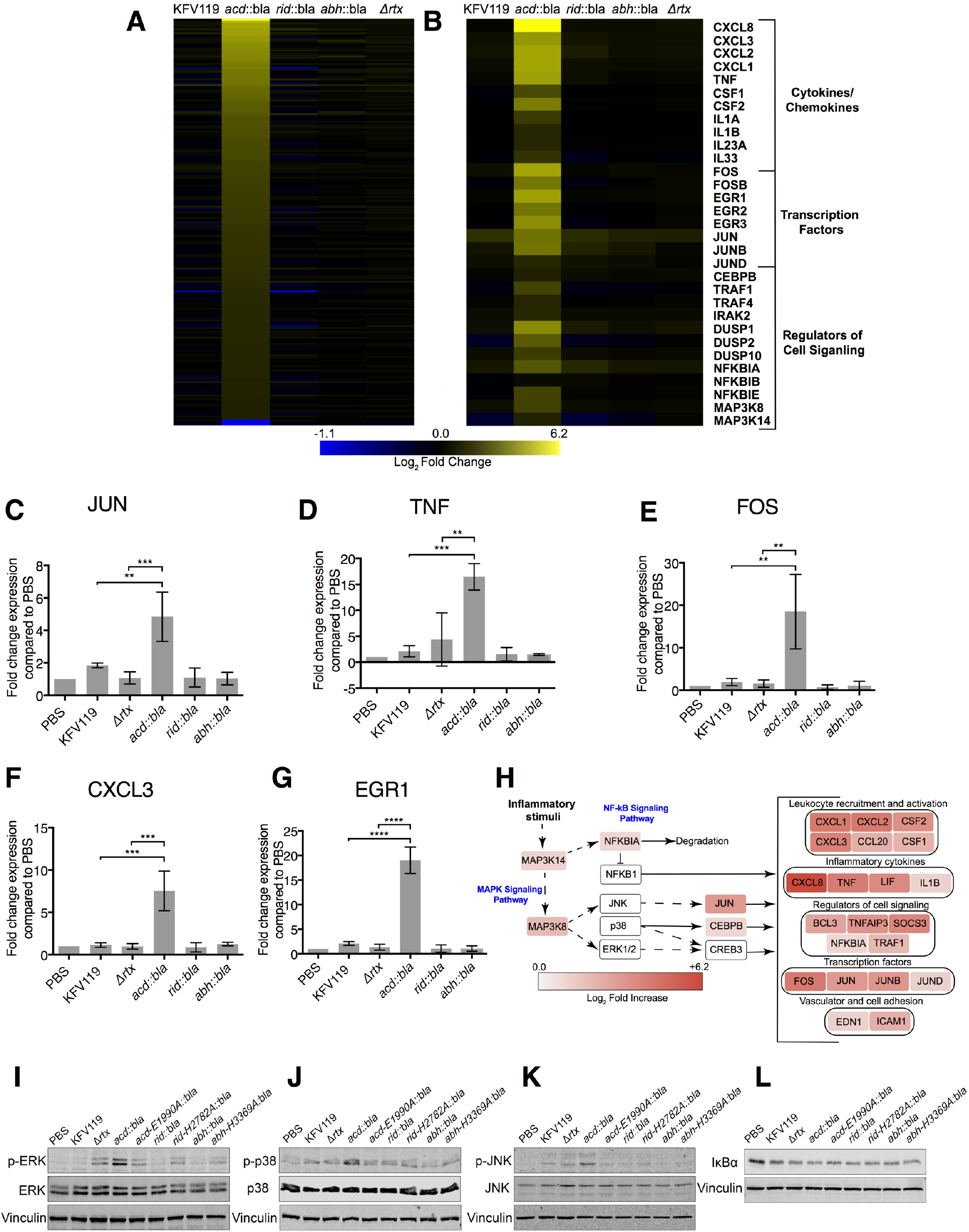
ACD induces a global proinflammatory response that is silenced when delivered with RID and ABH on the wild-type toxin. (**A, B**) Heat maps of differentially expressed genes identified in RNA-sequencing analysis of T84 cells inoculated with *V. cholerae* strains as indicated. Heat maps represent statistically significant differentially expressed genes with a −1 ≤ 0 ≤ 1 log_2_ fold change cut off. *n* = 2 independent experiments. (**C-G**) qPCR validation of select differentially expressed genes, *n* = 3 independent experiments. Data are reported as the mean ± s.d. (** p<0.01, ***p<0.001, ****p<0.0001, One-way ANOVA with Tukey’s multiple comparison’s test for samples as indicated. No other strains induced a statistically different response compared to the normalized PBS control or another strain). (**H**) ACD differentially regulated genes involved in regulation or downstream of the MAPK and NF-κB signaling pathways as identified from RNA-seq analysis. (**I-L**) Western blot analysis of phosphorylated ERK (p-ERK, I), phosphorylated p38 (p-p38, J), phosphorylated JNK (p-JNK, K), and IκΒα degradation (L) in T84 cells inoculated with *V. cholerae* strain as indicated. Blots are representative of *n* =3 experiments with quantification shown in Fig. S2A-D.

Similarly, *rid::bla, abh::bla,* and *Δrtx* also did not induce significant changes in IEC gene expression by two hours. By contrast, the *acd::bla* strain induced differential regulation of over 200 genes (Table S2). Many of the significantly upregulated genes by ACD were identified as regulators of inflammation. These include genes for chemokines CXCL8 (IL-8) and CXCL3 (interleukin-3), the cytokine tumor necrosis factor (TNF), and for proinflammatory transcription factors JUN, FOS, and EGR1/2/3 (Fig 3B-G). Activation of select proinflammatory genes was validated by qPCR (Fig. 2H, 3C-G). These data suggest that when delivered independently, ACD induces proinflammatory gene expression more rapidly than by *V. cholerae* alone. However, when ACD is delivered on the complete MARTX*_Vc_* toxin, the response is abolished.

### RID suppresses host MAPK signaling via Rac1 to modulate the ACD proinflammatory response

Bioinformatic analysis of the *acd::bla* differentially regulated genes revealed that many were associated with the ERK, p38, JNK MAPK and NF-κB pathways (Fig. 3H). These pathways have also been associated with induction of IL-8 by purified *V. cholerae* flagellin (*29*). The phosphorylation of MAPK in bacterial-treated cells was monitored after two hours. While KFV119-treated cells showed no ERK phosphorylation, *Δrtx* activated ERK signaling (Fig. 3I). Further, *acd::bla* stimulated ERK phosphorylation above that of *Δrtx* alone (Fig. 3I, Fig. S2A). Both the *rid::bla* and *abh::bla* strains suppressed ERK phosphorylation, while *rid*-H2782A*::bla* and *abh-*H3369A::*bla* strains did not (Fig. 3I, Fig S2A). KFV119-treated cells also showed no phosphorylation of p38 or JNK. However, these MAPK pathways were activated by *acd::bla* (Fig. 3J and K, Fig. S2B and C). While previous studies suggest *V. cholerae* PAMPs may also activate these pathways (*29*), the observed limited or undetectable p38 and JNK MAPK phosphorylation by *Δrtx* alone is likely a result of timing, since *V. cholerae* does not induce expression of proinflammatory genes until six hours post-inoculation (Fig. 2I and 3A). Finally, none of the effector domains inhibited NF-κB, as measured by IκBα degradation (Fig. 3L, Fig. S2D).

Rac1 is a well characterized regulator of these pathways (*34, 35*). While KFV119 suppresses Rac1 activation through the biochemical activity of RID, *acd::bla* significantly induced active GTP-bound Rac1 when compared to both PBS and KFV119 treated cells (Fig. S1). In total, these data show that ACD should activate Rac1 to induce downstream MAPK. However, in the MARTX*_Vc_* holotoxin, direct inactivation of Rac1 by RID blocks this Rac1-dependent signaling and thus blocks all MAPK signaling.

### RID and ABH inhibit ACD induced MAPK signaling when stoichiometrically delivered from the same holotoxin

The lack of MAPK activation in MARTX*_Vc_*^+^ KFV119-treated cells suggests there is active suppression of ACD-induced MAPK signaling due to interplay between the MARTX*_Vc_* effector domains. However, these experiments rely on comparing strains with only one effector to those with either all or no effectors. Even though all Bla strains are known to deliver equivalent Bla activity (*2*), differences in IL-8 secretion across strains could result from minor variation in toxin expression, production, or effector delivery particularly during co-inoculation studies (Fig. 2G). To address this, six new strains were generated in which single codons were changed in *rtxA* to produce MARTX*_Vc_* toxins that retain natural processing and stoichiometrically equivalent delivery of all effectors, but with one, two, or all three of the effectors carrying point mutations in essential catalytic site residues (Fig. 4A and Fig. S3A-G). Thus, each experiment is internally controlled for toxin expression and natural effector delivery at 1:1:1. All ACD^+^ strains showed similar actin laddering patterns, indicative of equivalent ACD effector delivery (Fig. S4A and B).

**Fig. 4.**
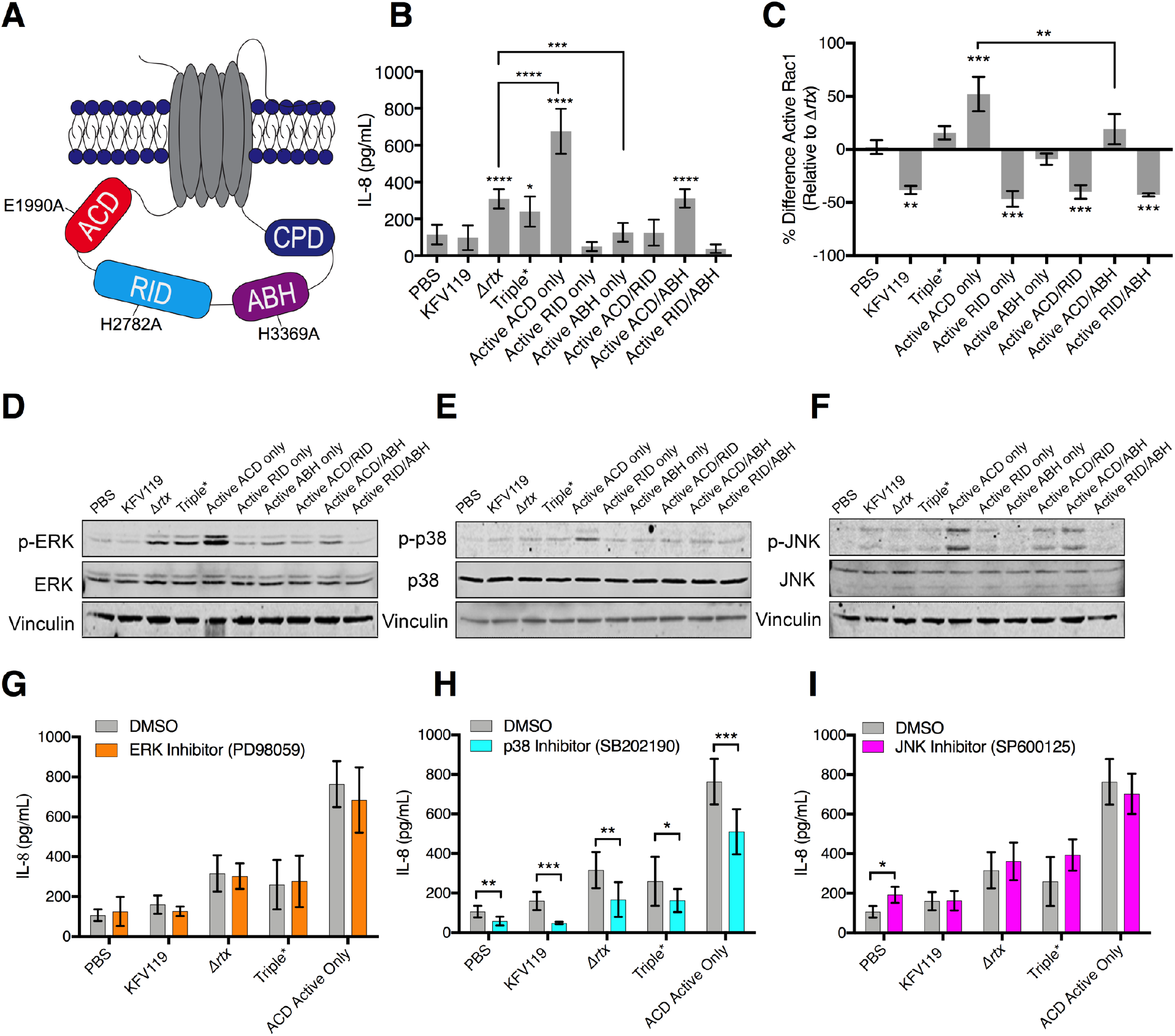
Both RID and ABH suppress ACD-induced proinflammatory responses. (**A**) Schematic of MARTX*_Vc_* toxin identifying catalytic residues mutated in single, double, and triple catalytically inactive MARTX*_Vc_* toxin effector strains. (**B, C**) Quantification of IL-8 secretion (**B**) or active (GTP-bound) Rac1 (**C**) measured from T84 cells inoculated with *V. cholerae* strains as indicated. For panel C, raw absorbances were normalized as the percent difference of active Rac1 relative to active Rac1 induced by the Δ*rtx* strain to identify effector specific changes. Data from *n* = 3 experiments reported as mean ± s.d. (*p<0.05, One-way ANOVA with Tukey’s multiple comparison’s test between cells treated with the indicated strain and PBS or for samples indicated, (** p<0.01, ***p<0.001, ****p<0.0001). (**D-F**) Western blot analysis of phosphorylated ERK(p-ERK, D), phosphorylated p38 (p-p38, E), and phosphorylated JNK (p-JNK, F) from T84 cells inoculated with strains as indicated. Blots are representative of *n* = 3 experiments. Quantification of data presented in Fig. S6. (**G-I**) Quantification of IL-8 secretion measured from T84 cells pre-treated with ERK inhibitor PD98059 (G), p38 inhibitor SB202190 (H), or JNK inhibitor SP600125 (I). Data pooled from *n* = 3 experiments reported as means ± s.d. (*p<0.05, ** p<0.01, ****p<0.0001, Student’s *t*-test for paired samples as indicated).

These strains were then tested for the impact of different effector domains activities on IL-8 secretion when simultaneously delivered from the same toxin. These studies confirmed results above with Bla strains that ACD alone induces IL-8 secretion, and that this activation is potently suppressed by RID (Fig. 4B). The amount of IL-8 secreted was equivalent between KFV119 and the ACD/RID active only strain demonstrating that RID dominates the inhibition of this pathway, independent of the presence of active ABH.

ABH also suppressed ACD-induced IL-8 secretion when introduced into cells from the same toxin (Fig. 4B) with inhibition varying across experiments from 20-50% (Fig. 4B and S5A-E). Thus, we considered that ABH might also inhibit Rac1 activation. As for the Bla strains, RID potently inhibited Rac1 activation compared to the Triple* strain and compared to the active ACD only strain (Fig. 4C). Although showing a lesser effect than RID, the ACD/ABH strain also reduced Rac1 activation compared to the active ACD only strain and this inactivation varied from 20-50% across independent experiments (Fig. 4C and 5S). The variable and more modest response supports that ABH inhibition of Rac1 is most likely indirect, which may explain the observed varying degree to which ABH suppresses both Rac1 activation and IL-8 secretion across independent experiments (Fig. 5SA-E); and why unlike RID, it does not completely abolish IL-8 secretion (Fig. 2F and 4B).

To confirm that both RID and ABH suppress ACD via MAPK signaling, activation of these pathways were examined in cells treated with the newly constructed strains. The single active ACD strain stimulated ERK activation beyond that of *V. cholerae* alone, while both the single active RID and ABH strains suppressed bacterial activation of ERK (Fig. 4D). Additionally, both the Active ACD/RID and Active ACD/ABH strain showed attenuation of ACD-induced ERK phosphorylation, JNK phosphorylation and p38 phosphorylation compared to the ACD Active only strain (Fig. 4D-F, Fig. S6A-C). Further, cells treated with PD98059, an inhibitor of ERK, showed no attenuation of the IL-8 response (Fig. 4G). However, inhibition of the p38 MAPK pathway by SB202190 did inhibit IL-8 secretion (Fig. 4H), while inhibition of the JNK pathway by SP600125 had no effect on IL-8 secretion (Fig. 4I).

In total, these data show that RID and ABH suppression of Rac1 and MAPK signaling pathways, particularly the p38 pathway, can modulate the ACD-induced proinflammatory response, and this response is attenuated by the action of RID and ABH.

### Cytoskeletal collapse activates proinflammatory signaling in ACD-treated cells in the absence of RID and ABH

ACD has two putative cytotoxic functions. The first is the inhibitor action of toxic actin oligomers (dimers and trimers) formed when only 2-4% of total actin has been cross-linked (*19, 36*). The second is ACD will eventually cross-link nearly 100% of cellular actin into higher order oligomers (10 to 15-mers) to sequester bulk actin and induce cytoskeletal collapse (*16*). Kinetic experiments comparing actin cross-linking abundance to MAPK activation were performed to determine which of these functions activate MAPK signaling. The ACD Active Only strain induced a maximum of 4% of total actin cross-linked between 5 and 15 min after addition of *V. cholerae*, corresponding to the formation of early toxic actin oligomers (Fig. 5A and B and Fig. S7A – C). Cross-linking increased around 60 min (Fig. 5A and B) corresponded to ACD sequestering bulk actin to induce actin depolymerization and cytoskeletal collapse. In the same assayed samples, MAPK signaling was activated between 60 and 90 min (Fig. 5C-H, Fig. S8A-B), when over 50% total monomeric actin was cross-linked. These results indicate that a significant portion of actin needs to be cross-linked to activate proinflammatory signaling. While the Triple* strain showed stochastic activation of MAPK signaling between 5 and 15 min, it induced significantly less ERK phosphorylation than the ACD Active Only strain and there was no detection of phosphorylated p38 or phosphorylated JNK between 30 and 120 min (Fig. S8C-H). This stochastic activation by the bacteria alone could account for the transient activation of p38 phosphorylation at 10 min that was observed in one experiment (Fig. 8A and 8B). These data suggest that it is ACD sequestration of bulk actin and cytoskeletal destruction that leads to activation of proinflammatory signaling in IECs.

**Fig. 5.**
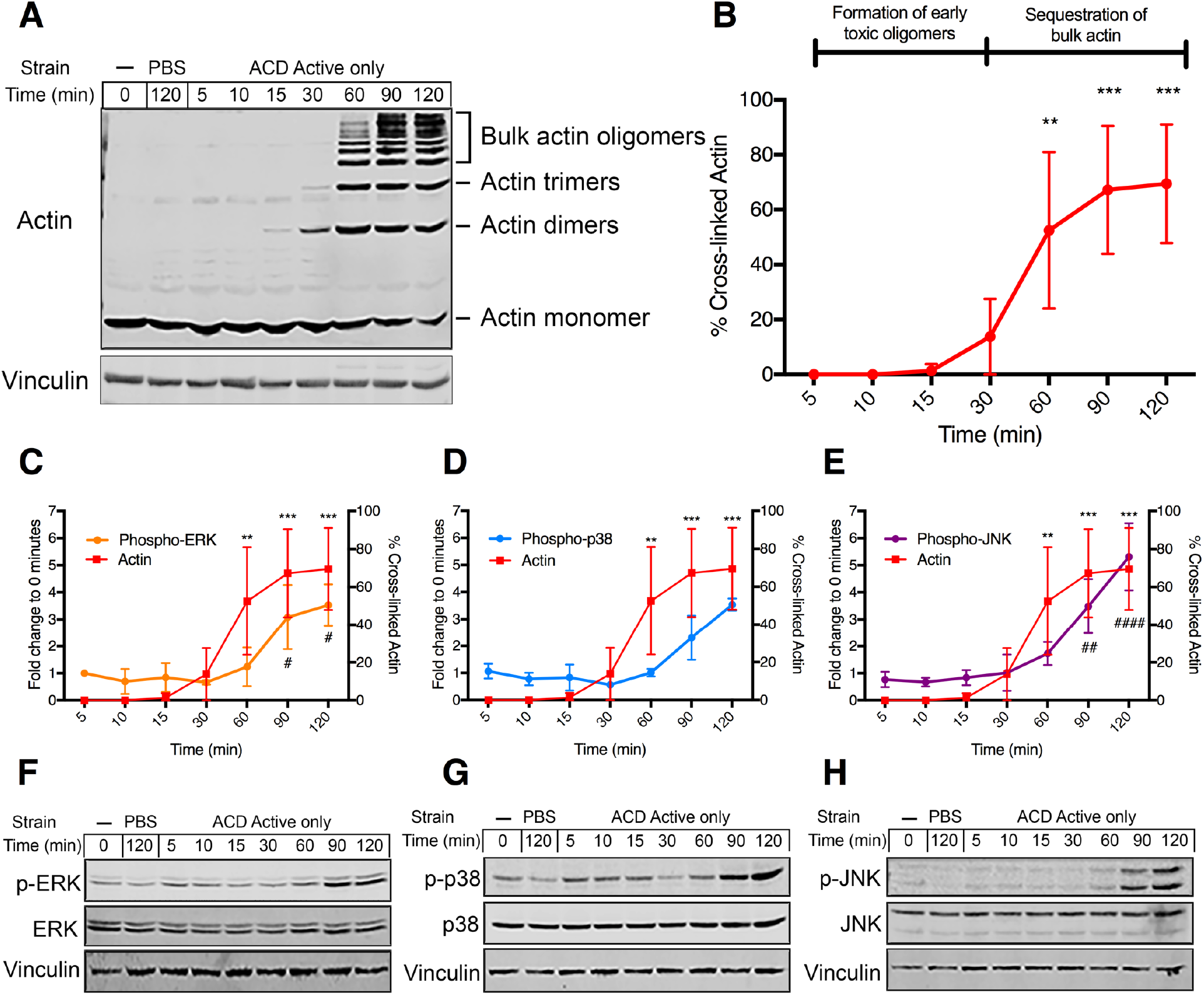
ACD activates proinflammatory MAPK signaling following sequestration of bulk actin. (**A, B**) Representative western blot and quantification (*n*=3 independent experiments) of actin cross-linking from T84 cells inoculated with the ACD only strain of *V. cholerae*. Data are reported as means ± s.d. (** p<0.01, ***p<0.001, One-way ANOVA with Tukey’s multiple comparison’s test between indicated timepoints and 0% actin cross-linking from 0 min control). (**C-E**) Fold change and (**F-H**) representative western blot of phosphorylated ERK (p-ERK, C), p38 (p-p38, D), or JNK (pJNK, E). All samples assayed from same experiment so actin crosslinking is duplicated across panels B-E. Data are reported as means ± s.d. (## p<0.01, ### p<0.001, #### p<0.0001, One-way ANOVA with Tukey’s multiple comparison’s test between phosphorylated ERK/JNK at indicated timepoints and a 0 min control with a normalized fold change of 1, *n* = 4 (C), 2 (D), or 3 (E) independent experiments. A third biological replicate was performed for phospho-p38 quantification but the ACD Active only strain induced a stochastically high 273-fold change compared to 0 min control (Fig. S6A-B).

To completely block the intestinal inflammatory response, RID and ABH would have to inactivate MAPK signaling prior to host detection of ACD-induced cytoskeletal collapse. This would indicate that instead of reversing the activation state of MAPK pathways after 60-90 min, RID and ABH inactive these pathways ahead of ACD-induced activation. Cells treated with KFV119 failed to induce any changes in MAPK activation following significant actin cross-linking (Fig. 6A-F). Further, both the ACD/RID Active strain and the ACD/ABH Active strain showed reduced phosphorylation of ERK, p38, and JNK, even following significant actin cross-linking at 60 min (Fig. S9A-L). Additionally, MAPK activation induced by the Active ACD/RID and Active ACD/ABH strains was still significantly less than activation by the ACD Active only strain at 120 min (Fig. 6G-I). These data would suggest that RID and ABH are each sufficient to attenuate ACD induction of MAPK signaling. However, modest, yet significant, activation of the JNK pathway by the Active ACD/RID strain was still observed at 120 min (Fig. S9A-F). The Active ACD/ABH strain also induced slight, yet significant, activation of the p38 and JNK pathways at 120 min (Fig. S9G-L). Since both RID and ABH alone do not completely abolish activation of these pathways, both effectors may be required to completely abolish the global inflammatory response observed in the RNA-seq analysis (Fig. 3A). The inhibition of MAPK signaling was not due to RID and ABH directly inhibiting ACD actin cross-linking activity. In fact, the ACD/ABH strain showed slight, yet significant, increase in actin cross-linking at 30 and 60 min before returning to similar abundances at 90 and 120 min (Fig. S9M). Therefore, both RID and ABH independently silence signal transduction pathways prior to host detection of ACD induced collapse of the actin cytoskeleton without modulating ACD activity.

**Fig. 6.**
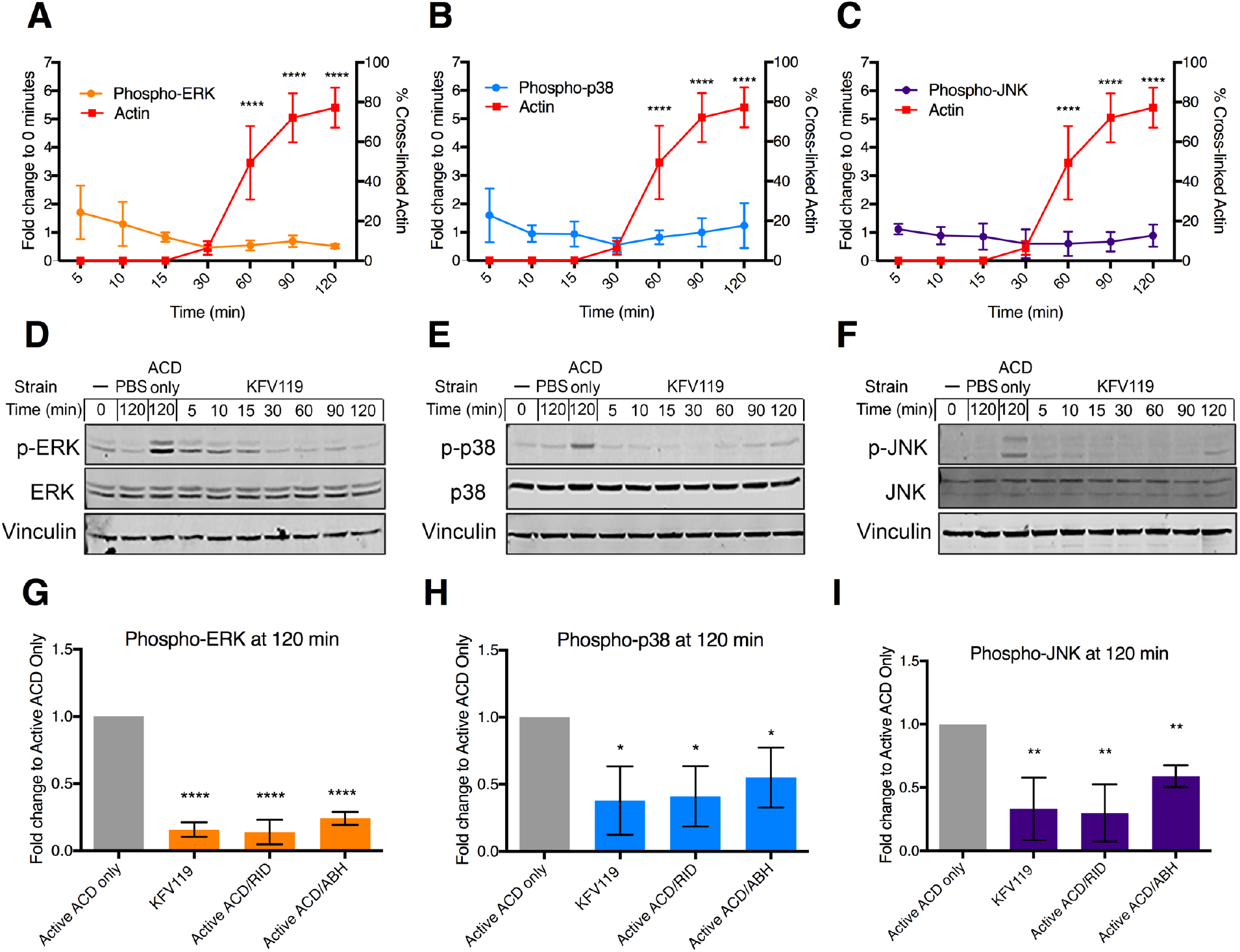
RID and ABH block MAPK signaling prior to host detection to ACD sequestration of bulk actin. (**A-C**) Fold change and (**D-F**) representative western blot of phosphorylated ERK (p-ERK, A), p38 (p-p38, B), or JNK (pJNK, C) from cells treated with *V. cholerae* KFV119. All samples assayed from same experiment so actin crosslinking is duplicated across panels A-C. Data from densitometry of *n* = 3 western blots reported as the mean ± s.d. (****p<0.0001, One-way ANOVA with Tukey’s multiple comparison’s test between indicated timepoints and 0% actin cross-linking from 0 min control). (**G-I**) Quantification of fold change in phosphorylated ERK (p-ERK, **G**), phosphorylated p38 (p-p38, **H**), and phosphorylated JNK (p-JNK, **I**) of samples as indicated compared to the Active ACD only strain at 120 minutes post-inoculation. Data from densitometry of *n* = 3 western blots reported as the mean ± s.d. (*p<0.05, **p<0.01, ****p<0.0005, Student’s *t*-tests compared to Active ACD only strain). No statistically significant difference was observed between KFV119, ACD/RID Active, or ACD/ABH Active strains compared to each other.

### RID can inhibit IL-8 secretion due to actin destruction by ACD and by latrunculin A, independent of live bacteria or formin inhibition

Between RID and ABH, RID is the more potent inhibitor of ACD-induced inflammatory response (Figs. 2F, 2G, 4B, and 4C). To determine if the action of RID is specific to MARTX toxins or more broadly applicable, cells were treated with purified recombinant RID or ACD fused to the N-terminal of the anthrax lethal toxin (LF_N_RID and LF_N_ACD, respectively) in combination with the anthrax protective antigen (PA). This previously characterized system allows for the delivery of MARTX effector domains into cells in the absence of bacteria and MARTX toxin delivery (Fig. 7A) (*37*). LF_N_RID has been demonstrated to acylate Rac1 in cells (*20*) and LF_N_ACD is sufficient for actin crosslinking (*37*). Cells treated with LF_N_ACD and PA induced IL-8 secretion, demonstrating that ACD is sufficient for induction of a proinflammatory immune response in the absence of a bacterium (Fig. 7B). The effects were not limited to the toxic action of ACD. Indeed, cells treated with latrunculin A, a sponge toxin that binds and sequesters G-actin, also showed IL-8 secretion. Treatment of cells with LF_N_RID in the presence of PA suppressed latrunculin A induced IL-8 secretion (Fig. 7C). However, inhibiting formins with SMIFH2 failed to elicit an IL-8 response (Fig. 7D). These data support that it is the onset of cytoskeletal collapse, rather than limited cytoskeletal damage as induced by SMIFH2 that stimulates the inflammatory pathways and, regardless of the inducer, these pathways are inhibited by RID.

**Fig. 7.**
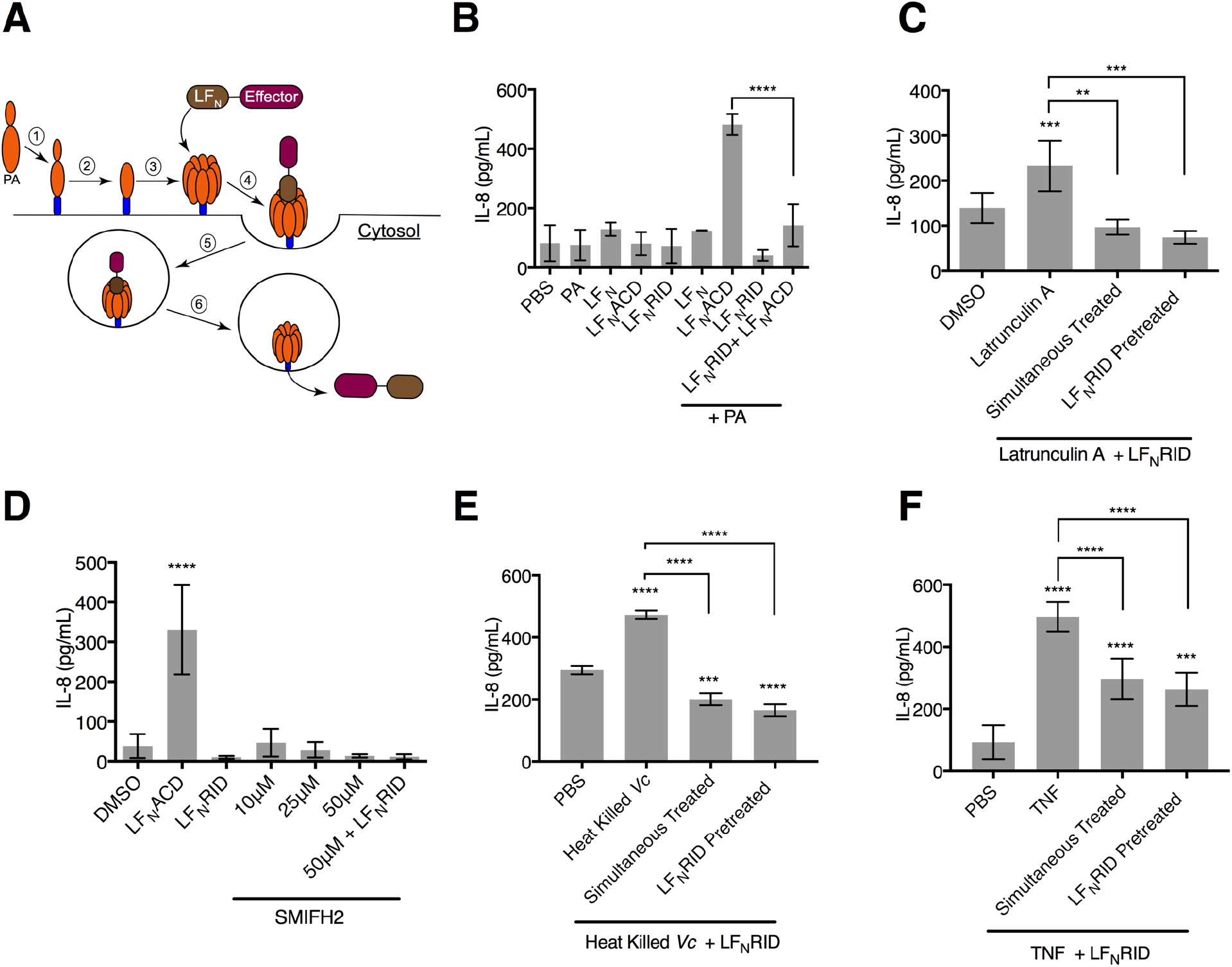
Cytoskeletal collapse, and not formation of toxic actin oligomers, activates proinflammatory response in IECs. (**A**) Schematic of LF_N_-Effector intoxication. (1) PA binds to the anthrax toxin receptor. (2) PA is processed to its 63 kDa active form (PA_63_). (3) PA_63_ oligomerizes to a heptamer complex. (4) The LF_N_ domain of the LF_N_-MARTX effector fusion binds to the PA_63_ heptamer. (5) The LF_N_-Effector + PA63 complex enters the cell through receptor mediated endocytosis. (6) Acidification of the vacuole promotes the PA_63_ heptamer to form a pore in the vacuole membrane to release LF_N_-Effector fusion protein into the cytosol. (**B**) Quantification of IL-8 secretion measured from T84 IECs treated with LF_N_ alone, LF_N_ACD, or LF_N_RID in the presence or absence of PA. Data are from *n* = 3 biological replicates reported as means ± s.d. (****p<0.0001, One-way ANOVA with Tukey’s multiple comparisons). (**C, D**) Quantification of IL-8 measured from T84 cells treated with **(C)** latrunculin A or SMIFH2 alone or in combination with LF_N_RID in the presence of PA where indicated. Data are *n*= 6 biological replicates reported as means ± s.d. (*** p<0.001, ****p<0.0001, One-way ANOVA with Tukey’s multiple comparison’s test). (**E, F**) Quantification of IL-8 measured from T84 cells treated with heat-killed *V. cholerae* (*Vc*) or (**F**) TNF in combination with LF_N_RID in the presence of PA where indicated. Data from *n*=3 (**E**) or *n*=6 (**F**) biological replicates reported as means ± s.d. (**p<0.01 *** p<0.001, ****p<0.0001, One-way ANOVA with Tukey’s multiple comparison’s test compared to PBS only or for samples as indicated).

**Fig. 8.**
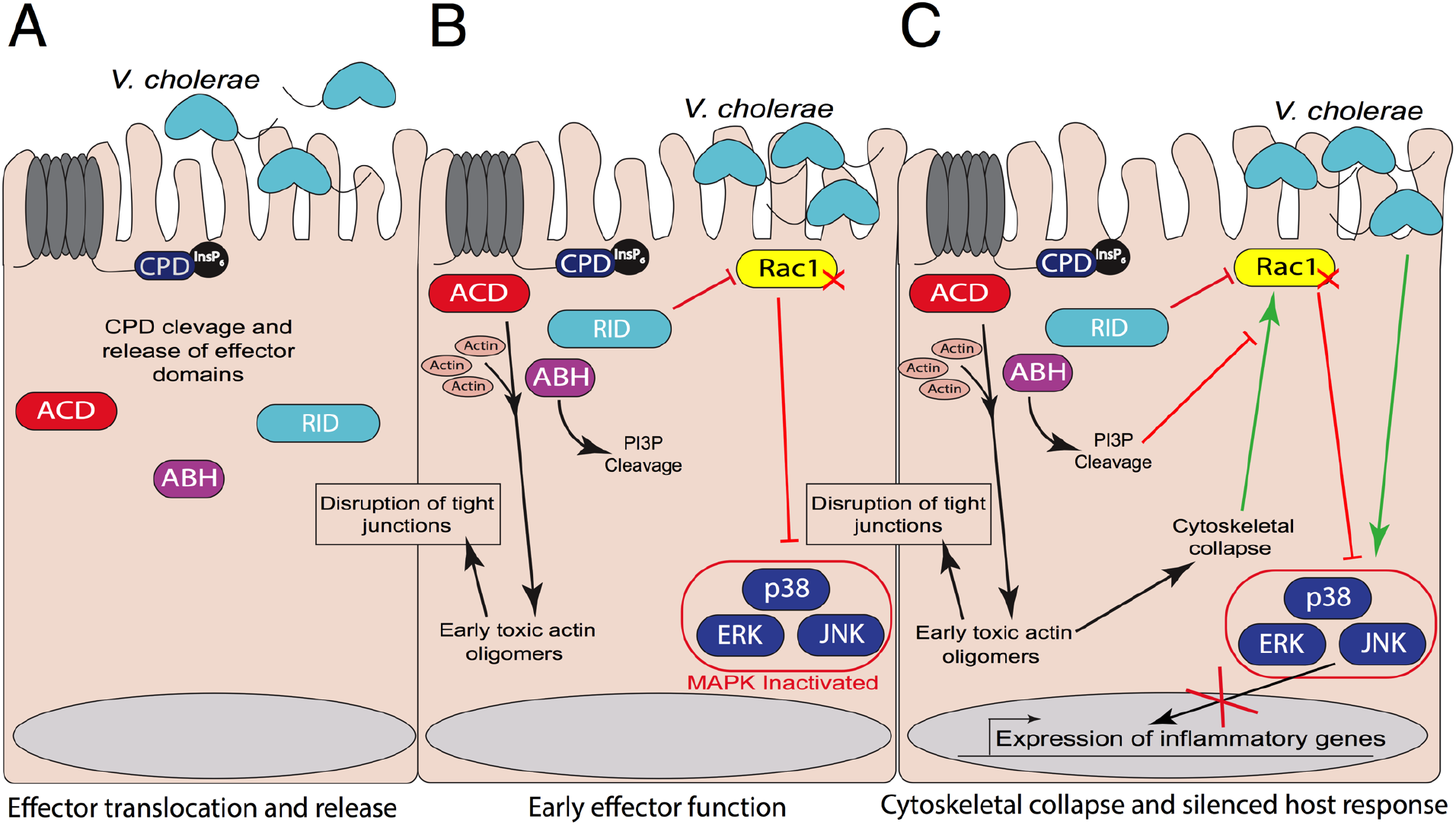
Model of interplay between the MARTX*_Vc_* toxin multiple functions. (**A**) MARTX*_Vc_* toxin effector domains are separated from the holotoxin. (**B**) During the early stages of effector activity, ACD actin cross-linking forms toxic actin oligomers to disrupt tight junctions. ABH cleaves PI3P and RID inactivates Rac1 to block activation of MAPK signaling pathways. (**C**). Inactivation of Rac1 and its upstream signaling pathways prevent MAPK activation to silence the inflammatory response to ACD destruction of the cytoskeleton and host detection of bacterial PAMPs.

This inhibition of proinflammatory immune response to bacterial factors was also more broadly applicable. Treatment of cells with LF_N_RID and PA was also sufficient to suppress IL-8 secretion induced by heat-killed bacteria (Fig. 7E) and to attenuate IL-8 secretion in response to purified TNF (Fig. 7F). The residual stimulation of IL-8 by TNF most likely occurs through TNF induction of NF-κB signaling, which RID does not inhibit (Fig. 3L).

All total, these data demonstrate that RID is a potent inhibitor of inflammation induced by PAMPs and by distinct pathways of cytoskeletal damage, indicating RID is also a broadly applicable inhibitor of damage-associated molecular patterns (DAMPs) (Fig. 8A-C).

## DISCUSSION

MARTX toxins are unique hybrids of independently secreted toxins and multifunctional effector delivery systems (*8, 38*). This study demonstrates that the *V. cholerae* MARTX toxin utilizes this multifunctionality to “self-regulate” and silence the host response both to its own cytotoxic activity and detection of bacterial PAMPs. Following translocation, ACD begins cross-linking actin to produce early toxic oligomers that sequester actin binding proteins and disrupt intestinal tight junctions. While these toxic oligomers are being formed, RID directly inactivates Rac1 and ABH cleaves PI3P to directly or indirectly regulate Rac1 activation. At this early timepoint, the downstream MAPK signaling pathways are silenced. Once ACD begins to sequester bulk actin to induce cytoskeletal collapse, the cell is unable to trigger Rac1 and MAPK activation. Therefore, the MARTX*_Vc_* toxin blocks IECs from activating an innate immune response (Fig. 8A-C). While previous studies have suggested MARTX^+^ *V. cholerae* strains induce IL-8 secretion, those studies found this proinflammatory response to be growth phase dependent in which stationary phase cultures stimulated more IL-8 release than log phase cultures (*27*). Secretion of the MARTX*_Vc_* toxin is also growth phase dependent and is expressed and secreted during log phase and then toxin present in the supernatant fluids is degraded by proteases secreted during stationary phase (*39*). Therefore, no toxin would be present to suppress IL-8 secretion in stationary phase culture fluids. Our findings support a new model in which MARTX*_Vc_* toxin produced during the early active growth stage in the intestines acts to suppress the IEC inflammatory response. These data indicate that MARTX*_Vc_* toxin suppression of intestinal innate immunity prevents host recruitment of immune cells to protect the colonizing bacteria from neutrophil-mediated clearance.

Additionally, MARTX*_Vc_* toxin immunomodulatory activities may contribute to the differences in inflammation observed between various *V. cholerae* strains (*27, 28*). In particular, the current predominant circulating “altered” El Tor *V. cholerae* strains responsible for pandemic disease since the mid 2000’s are hypervirulent and exhibit increased clinical severity of diarrhea (*40, 41*). This is in part due to the acquisition of mutations in H-NS and VieA resulting in increased production of cholera toxin and HlyA, increased motility, and inflammasome activation suggesting increased inflammation (*33, 41, 42*). In addition, these strains naturally lack the MARTX*_Vc_* toxin due to a stop codon in *rtxA* (*15*). Restoring toxin secretion in altered El Tor isolate 2010EL-1786 attenuated IL-8 induction comparable to the parent strain (Fig. 1C). Thus, loss of the MARTX*_Vc_* toxin immunomodulatory activities in these altered strains may increase intestinal inflammation, which could exacerbate disease severity and contribute to its hypervirulence.

The effector domains, and not the MARTX*_Vc_* pore, were identified to modulate innate immune signaling in IECs. These data support a previously established model in which the MARTX pore functions primarily as an effector delivery platform and not a direct virulence mechanism (*43*). How each delivered effector contributes to pathogenesis is still debated. While other studies have connected RID and ABH to inhibition of macrophage phagocytosis in vitro (*20, 44*), these conclusions contradict previous findings that 5% inhibition by these effectors is biologically insignificant compared to the 90% inhibition by ACD (*2*). Our study validates that RID and ABH most likely function primarily to abolish host detection of cytoskeletal damage and bacterial PAMPs. RID-mediated inactivation of MAPK pathways most likely occurs through its direct inactivation of Rac1 and direct or indirect inactivation other Rho family GTPases, thereby blocking ACD upregulation of these GTPases. Rho GTPases regulate MAPK and other cell signaling pathways and bacterial toxin inactivation of Rho GTPase can inhibit these pathways in specific cell types (*34, 35*). However, toxin inactivation of Rho GTPases through covalent modifications, such as ampylation or glycosylation, induces IL-8 secretion in IECs and activates the pyrin inflammasome in macrophages (*45, 46*). However, RID does not activate the inflammasome (*46*). Therefore, RID acylation of the C-terminal polybasic region of Rho family GTPases allows for the inactivation of Rho GTPases to suppress proinflammatory signaling pathways in IECs, while also evading host detection of Rho inactivation through the inflammasome.

How cleavage of PI3P by ABH blocks Rac1 activation and proinflammatory signaling is still unknown. Some Rho family GTPases utilize guanine nucleotide exchange factors (GEFs) whose activity is regulated by phosphatidylinositol binding to promote activation of Rho GTPases (*47*). For example, the C-terminal domain of the Rac1 GEF Tiam-1 has weak binding affinity to PI3P. While this activity is not required for Tiam-1 subcellular localization, abolishing Tiam-1 PI3P binding inhibits Rac1 activation (*48*). ABH depletion of PI3P may also disrupt the equilibrium of other phosphatidylinositol lipids, such phosphatidylinositol (3,4,5) triphosphate (PIP3) or phosphatidylinositol (3, 4) diphosphate (PIP2), which spatially regulate Rac1 GEFs and GTPases activating proteins (GAPs) and control formation of active Rac1 nanoclusters in membranes (*49, 50*). Additionally, PI3P is required for the formation of autophagosomes and ABH-mediated cleavage of PI3P inhibits formation of autophagosomes and endocytic trafficking (*22*). ABH depletion of PI3P could also block formation of scaffolding complexes on autophagosomes and endosomes that promote recruitment and activation of MAPK signaling molecules (*51, 52*). The loss of the scaffolds would further prevent activation of the MAPK pathway.

ACD production of toxic actin oligomers has been characterized as the primary cytotoxic mechanism of ACD based on the hypothesis that the kinetics of ACD actin cross-linking prevent it from cross-linking the majority of cellular actin (*19*). Our study reveals ACD stimulation of MAPK signaling pathways occur following cross-linking of over 50% of cellular actin (Fig. 6, 7). This supports previous findings in which type VI secreted ACD cross-links significant quantities of actin and induce intestinal inflammation in mice (*53*). While the toxic oligomers may still contribute to cytoskeletal collapse, our data indicate that it is the destruction of the cytoskeleton, and not the direct formation of toxic actin oligomers inhibition of formins, that activate proinflammatory signaling.

Synergistic and antagonistic interactions between co-delivered bacterial effectors can have a profound impact on the host response and virulence. Type III secreted effectors SopE and SopB from *Salmonella enterica* serovar Typhimurium work in synergy to recruit MYO6 to the plasma membrane to facilitate actin rearrangement and promote bacterial invasion (*54*). Other combinations of *S. enterica* Type III secretion effectors have also been shown to display synergistic and antagonistic interactions (*55*). YopE and YopT from *Yersinia pestis* independently activate the inflammasome. However, inflammasome activation is suppressed during infection by the co-delivered effector YopM and *Y. pestis* strains lacking YopM are less virulent in mice due to the immune response from the inflammasome (*56*). Similarly, RID and ABH silencing the global response to ACD activity provide evidence that different MARTX effector domain combinations could impact the overall host response to co-delivered effectors. Effector translocation from the same toxin assures stoichiometric delivery that can further maximize the synergistic benefit. These data suggest that an effector’s contribution to virulence and disease could be enhanced or attenuated depending on which other effector domains it is delivered with. *Vibrio vulnificus* with different natural effector combinations have varying virulence potential and promote altered host responses (*57–59*). Therefore, MARTX toxin multifunctionality, in combination with variability in the effector domain repertoire, allows for variety of effector interplay combinations which could either enhance or attenuate MARTX toxin associated virulence.

## MATERIALS AND METHODS

### Antibodies and chemical reagents

Antibodies used in this study include phospho-p44/p42 MAPK (ERK1/2) (Cell Signaling Technology, #4337S), p44/p42 MAPK (ERK1/2) (Cell Signaling Technology, #4965S), phospho-p38 MAPK (Cell Signaling Technology, #9211S), p38 MAPK (Cell Signaling Technology, #9212S), phospho-SAPK/JNK (Cell Signaling Technology, #9255S), SAPK/JNK (Cell Signaling Technology, #9252S), Vinculin (Cell Signaling Technology, #13901), IKBα (Cell Signaling Technology, #9242S), Tubulin (Cell Signaling Technology, #2144), GAPDH (Santa Cruz, #sc-25779), Actin (Sigma, #A2066), and LICOR IRDye 800CW/680LT secondary antibodies (LICOR, #926-3221, #926-3211, #926-68070, #926-68071). Chemical reagents used in this study were ERK MAPK inhibitor PD98059 (Cell Signaling Technology, #9900L), p38 MAPK inhibitor SB202190 (Cell Signaling Technology, #S7067), JNK MAPK inhibitor SP600125 (Abcam, #120065), Latrunculin A (Millipore, #428021), and SMIFH2 (Fisher Scientific, #440110) and recombinant human TNF (R&D Systems, #210-TA-005/CF).

### Cell culture

T84 male colorectal carcinoma cells acquired from the American Type Culture Collection (ATCC, #CCL-248) were cultured in Dulbecco’s Modified Eagle Medium/Nutrient Mixture F-12 (DMEM/F12), GlutaMAX supplement (ThermoFisher Gibco, #10565018) with 10% (w/v) fetal bovine serum (FBS, Gemini Bio-products, #900-108) and 0.1% penicillin/streptomycin (ThermoFisher Gibco). Hela cells were grown in DMEM (ThermoFisher Gibco, #11965) with 10% FBS and 0.1% penicillin/streptomycin. Cells were maintained at 37°C in the presence of 5% CO_2_.

### Bacterial growth medium

Bacterial strains and plasmids used in this study are listed in Table S3. All *V. cholerae* strains used in this study are derived from spontaneous streptomycin-resistant derivatives of clinical isolates N16961 or 2010EL-1796. *V. cholerae* and *Escherichia coli* were grown on Luria-Bertani (LB) broth or agar. *V. cholerae* medium was supplemented with 100 μg mL^-1^ streptomycin, 2 μg mL^-1^ chloramphenicol, 100 μg mL^-1^ ampicillin, or 5% (w/v) sucrose as needed. *E. coli* growth medium was supplemented with 10 μg mL^-1^ chloramphenicol, 100 μg mL^-1^ ampicillin, or 50 μg mL^-1^ kanamycin as needed.

### Treatment of cells with live bacteria

*V. cholerae* strains were grown at 30°C in LB medium supplemented with 100 μg mL^-1^ streptomycin. Overnight cultures were diluted 1:100 and grown at 30°C with shaking until exponential phase (OD_600_≍0.40-0.60). Bacteria from 1 mL were pelleted by centrifugation and resuspended in phosphate buffered saline (PBS) to a final concentration of 5 x 10^8^ bacterial cells mL^-1^. T84 cells (seeded the day before into cell culture-treated plates) were twice washed with PBS and then media changed to antibiotic- and FBS-free DMEM/F12. Resuspended *V. cholerae* was added to media over cells (multiplicity of infection = 5). Inoculations were synchronized by centrifugation at 500 x*g* for 3 min. T84 cells were subsequently incubated at 37°C in the presence of 5% CO_2_ until processed for downstream applications.

### LF_N_ Effector intoxication of T84 cells

LF_N_ fusion proteins were expressed from pTCO24 (LF_N_ACD) and pKS119 (LF_N_RID) and purified as previously described (*21, 37*). In brief, *E. coli* containing BL21(DE3)(pMagic) with overexpression plasmids were grown in Terrific broth and expression of protein induced with 1 mM isopropyl β-D-thiogalactopyranoside at 25°C overnight. Bacteria were harvested in Buffer A (10mM Tris, 500 mM NaCl pH 8.3), lysed by sonication, and lysate clarified by centrifugation at 16,0000 x*g* for 30 min. The 6xHis-tagged recombinant proteins were purified using Ni-NTA HisTrap column followed by size-exclusion chromatography using a Superdex 75 column in Buffer A with 5 mM β-mercaptoethanol using the ÄKTA protein purification system (GE Healthcare). Proteins were stored in 10% glycerol at −80°C.

For intoxication of cells, media was replaced over 10^5^ cells previously seeded in a 12-well plate. 31.7 nM PA alone (List Labs, #171E) or in the presence of 13.6 nM LF_N_, LF_N_-ACD, or LF_N_-RID was added to media and cells were incubated for 20 hr at 37°C in the presence of 5% CO_2._ For co-intoxication of 13.6 nM LF_N_ACD and LF_N_-RID, 3x excess PA was used to ensure equal translocation of both effectors. Co-treatment of 1μM latrunculin A or 5 ng/mL of recombinant TNF were used when indicated. For studies using co-treatment of boiled bacteria, 1 mL of an overnight culture of the Δ*rtx* was pelleted by centrifugation and resuspended in 1 mL of PBS. The resuspended bacteria were boiled for 10 minutes at 95°C. 20 μL of the boiled culture was used to stimulate IL-8 secretion in T84 cells were indicated.

### Construction of pDS132 *sacB*-counterselectable plasmids

Fragments (gBlocks) containing either the ACD E1990A mutation or the ABH H3369A mutation along with 500 base pairs up and downstream from the mutations were commercially synthesized. Each fragment also has additional secondary silent mutations to introduce novel MfeI and PstI restriction sites near the E1990A and H3369A mutations, respectively. Sequences are listed in Table S4. Fragments were cloned into pDS132 digested with SphI (New England Biolabs, #R3182S) using Gibson Assembly (New England Biolabs, #E2611S) according to manufacturer’s instructions. Plasmids were recovered and propagated in DH5αλ*pir* on LB supplemented with chloramphenicol and confirmed by sequencing.

### Transfer of E1990A, H2782A, and H3369A mutations to the *V. cholerae* chromosome

The ACD E1990A and ABH H3369A mutations on the above pDS132-based plasmids were transferred to SM10λ*pir*. These mutations or the RID H2782A mutation from pSA129 were transferred to the *V. cholerae* chromosome by conjugation followed by *sacB*-dependent counterselection for double homologous recombination as previously described (*2, 60*). To confirm recombinants gained the desired mutations, regions corresponding to the mutations were amplified by PCR, digested with the introduced novel restriction site, and products were separated on agarose gel. Presence of introduced mutations were also confirmed by sequencing.

### IL-8 enzyme-linked immunosorbent assay (ELISA)

1- to 2 x 10^5^ T84 cells seeded in a 12-well tissue culture treated plate were treated as described above for 2 hr. Media from inoculated cells was removed, cells were washed once with warm PBS and media changed to DMEM12 GlutaMAX media supplemented with serum, pen-strep, and 100 μg mL^-1^ gentamicin. Cells were incubated for an additional 20 hr at 37°C in the presence of 5% CO_2_. For IL-8 chemical inhibitor studies, 10 μM of MAPK inhibitor or 0.33% DMSO control were added to T84 cells one hour prior to bacterial challenge. Inhibitors were reapplied following 2 hr bacterial challenge. Media from the T84 cells was harvested and spun down at 20,000x*g* for 1 min at 4°C and supernatant was collected. Concentration of IL-8 in cell media was measured using the IL-8 Human Matched Antibody Pair ELISA kit (ThermoFisher, #CHC1303) following manufacturer’s instructions.

### Western blot analysis of actin crosslinking and cell signaling pathways

1 x 10^6^ T84 cells in a 6-well tissue culture dish or 1 x 10^5^ Hela cells in a 12-well dish were treated with various *V. cholerae* strains as described above for 2 hr. Cells were washed once with cold PBS and removed from the plate in 150 μL lysis buffer (150 mM NaCl, 20 mM TRIS pH 7.5, 1% Triton X-100, and Pierce Protease and Phosphatase inhibitor added prior to use) using a cell scrapper. Lysates were incubated on ice for 15 min and then clarified by centrifugation at 20,000*xg* at 4°C for 10 min. Concentration of protein in the collected supernatant fluid determined using the bicinchoninic (BCA) assay (ThermoFisher, #23227). Specific samples were also collected directly in 150 μL 2X SDS loading buffer where indicated. Normalized samples were boiled for 5 min at 95°C in SDS loading buffer and protein separated on either a 10% or 15% SDS-polyacrylamide gel. Proteins were transferred to nitrocellulose membranes and blocked in TBS (10 mM Tris pH=7.4, 150 mM NaCl) with 5% (w/v) milk for 1 hour. Membranes were washed with TBS and then incubated in indicated primary antibodies 1:1000 in TBS with 5% (w/v) bovine serum album (BSA, Fisher Bioreagents) overnight at 4°C. Membranes were washed with TBS and probed in 1:10,000 IRDye 800CW/680LT secondary antibody for 1 hour before being washed again and imaged using the LI-COR Bioscience Odyssey imaging system. Quantification of band density was conducted using Fiji/ImageJ.

### Rac1 G-LISA

1 x 10^6^ T84 cells in a 6-well tissue culture dish were treated with various *V. cholerae* strains as described above for 2 hr. Cells were washed once with cold PBS and resuspended in 100 μL G-LISA lysis buffer. Activation of Rac1 was quantified from equal concentration of lysate using the Rac1 G-LISA Activation Assay (Cytoskeleton, #BK128) following manufacturers protocol. Percent change in active Rac1 compared to Δ*rtx* strain was calculated as (Abs_490_(Sample)-Abs_490_(Δ*rtx*))/ Abs_490_(Δ*rtx*))*100.

### RNA-seq

The stranded mRNA-seq was conducted in the Northwestern University NUSeq Core Facility. Briefly, total RNA examples were checked for quality using RINs generated from Agilent Bioanalyzer 2100. RNA quantity was determined with Qubit fluorometer. The Illumina TruSeq Stranded mRNA Library Preparation Kit was used to prepare sequencing libraries from 750 ng of high-quality RNA samples (RIN=10). The Kit procedure was performed without modifications. This procedure includes mRNA purification and fragmentation, cDNA synthesis, 3’ end adenylation, Illumina adapter ligation, library PCR amplification and validation. lllumina NextSeq 500 Sequencer was used to sequence the libraries with the production of single-end, 75 bp reads.

The quality of DNA reads, in fastq format, was evaluated using FastQC. Adapters were trimmed, and reads of poor quality or aligning to rRNA sequences were filtered. The cleaned reads were aligned to the human reference genome using STAR (*61*). Read counts for each gene were calculated using htseq-count (*62*). Normalization and differential expression were determined using DESeq2 (*63*). The cutoff for determining significantly differentially expressed genes was an FDR-adjusted p-value less than 0.05. A pathway analysis was performed on both gene lists using GeneCoDis (*64–66*) to identify pathways that are enriched with genes that are upregulated and downregulated.

### Quantitative RT-PCR

1 x 10^6^ T84 cells in a 6-well tissue culture treated plate were treated with *V. cholerae* as described above. At indicated time points, mRNA was harvested using the Qiagen RNeasy kit (Qiagen, #74014) following manufacturer’s instructions. RNA isolated was measured using a Nano-drop 100 spectrophotometer. Reverse transcription was performed using random hexamers (Roche) and Superscript III Reverse Transcriptase (Invitrogen, #18080044) in the presence of RNase OUT (Invitrogen, #10777019) or RNasin (Promega, #N2611) under the following conditions: 25°C for 5 min, then 55°C for 60 min, 95°C for 5 min. Remaining RNA was hydrolyzed using 1 N NaOH. Quantitative PCR was performed using iQ SYBR Green supermix (Bio-Rad, #1708880) on the iQ5 Multicolor RealTime PCR Detection System using gene specific primers indicated in Table S5. Relative change in gene expression compared to PBS control was determined the ΔΔCT method (*67*).

#### Quantification and statistical analysis

All experiments were done at least in triplicate and quantitative results are reported as the mean ± standard deviation (s.d.). Statistical analysis was performed using GraphPad Prism v6.0 as detailed in the figure legends. Statistical differences in ELISA and qPCR results were determined by one-way ANOVA followed by Tukey’s multiple comparison’s test. Statistical difference in results comparing suppression of ERK, p38, and JNK signaling by various *V. cholerae* strains compared to the ACD Active only strain were determined using Student’s t-tests.

## SUPPLEMENTARY MATERIALS

Fig. S1. ACD activation of Rac1 is abolished when delivered with RID on the wild type toxin.

Fig. S2. ACD induced phosphorylation of ERK, p38, and JNK is suppressed when co-delievered with RID and ABH.

Fig. S3. Schematic of single and double catalytically active MARTX*_Vc_* toxin effector stains and triple inactive MARTX*_Vc_* toxin effector strain.

Fig. S4. Actin laddering in T84 cells treated with *V. cholerae*.

Fig. S5. ABH mediated inhibition of host response to ACD varies across independent experiments.

Fig. S6. Both RID and ABH suppress ACD induced phosphorylation of MAPK pathways.

Fig. S7. Actin crosslinking in cells harvested in Triton X-100 lysis buffer.

Fig. S8. MAPK activation Triple* MARTX*_Vc_* toxin effector strain.

Fig. S9. RID and ABH block MAPK signaling without modulating ACD activity.

Table S1. MARTX effector-dependent transcriptional responses in *V*. *cholerae*-infected T84 cells.

Table S2. ACD-dependent transcriptional responses in in *V*. *cholerae*-infected T84 cells.

Table S3. Bacterial strains and plasmids used in study.

Table S4. Sequences of gBlocks used in the study.

Table S5. Primers used in study.

## Acknowledgements

We would like to thank the Northwestern University Center for Genetic Medicine NUSeq core facility, especially Xinkun Wang and Matthew Schipma, for technical assistance, bioinformatic analysis of the RNA-sequencing experiments and DNA sequencing. We would like to thank members of the Satchell lab for their valuable input and technical support and Dr. Nicholas Cianciotto and Dr. Gail Hecht for review of the manuscript.

## Funding

This work was supported by the NIH Ruth L. Kirschstein Institutional National Research Service Award Training Grant in Immunology and Microbial Pathogenesis T32AI007476 (to P.J.W.) and NIH grants R01AI092825 and R01AI098369 (to K.J.F.S.).

## Author contributions

P.J.W. conceptualized, designed, and conducted all experiments. K.J.F.S advised on all experiments. P.J.W. wrote the original draft of the manuscript and both P.J.W. and K.J.F.S. reviewed and edited the manuscript.

## Competing interests

The authors declare that they have no competing interests.

## Data and material availability

The RNA-seq data have been deposited in the NCBI Gene Expression Omnibus (GEO) under the accession number GSE125453. All other data needed to evaluate the conclusions in the paper are present in the paper or the Supplementary Materials.

