## Supplemental Figures and Tables 3 - 5 for "The *Vibrio cholerae* MARTX toxin simultaneously induces actin collapse while silencing the inflammatory response to cytoskeletal damage"

**SUPPLEMENTARY MATERIAL**

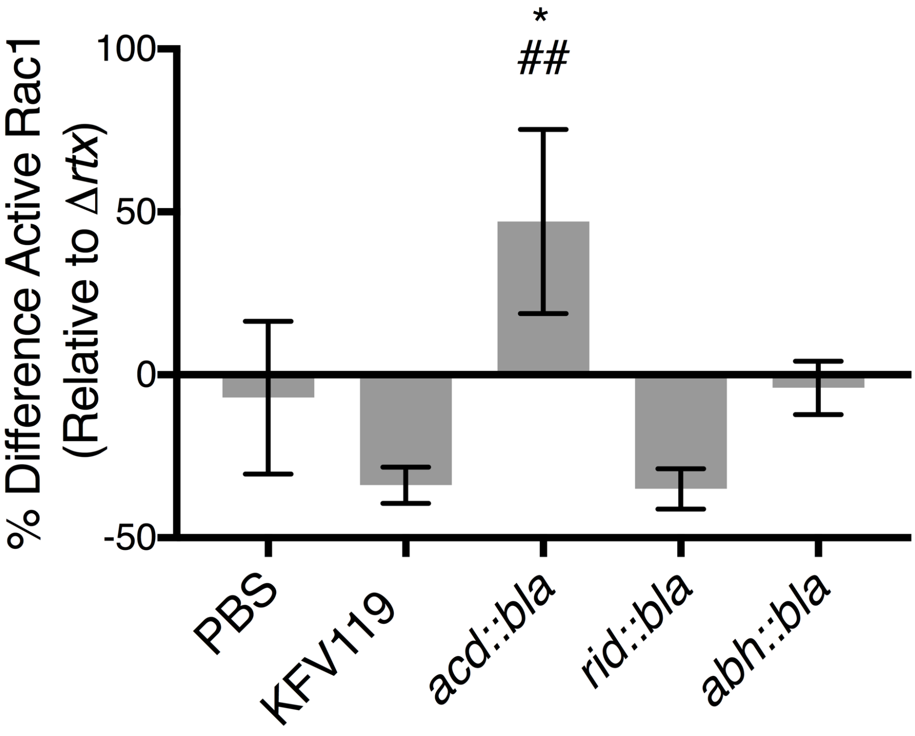


**Fig. S1. ACD activation of Rac1 is abolished when delivered with RID on the wild type toxin.** Active GTP-bound Rac1 was measured from cell lysates from T84 cells inoculated with various strains of *V. cholerae* by GLISA$^{TM}$. Raw absorbance data was normalized as the percent difference of active Rac1 relative to active Rac1 induced by the Δ*rtx* strain to identify effector specific, and not bacterial, activation of Rac1 at 120 minutes. *n* = 3 independent experiments. Data reported as mean ± s.d. (*p<0.05, One-way ANOVA with multiple comparisons between indicated T84 cells treated with the indicated strain and PBS. ##p<0.0, 1One-way ANOVA with Tukey's multiple comparisons between indicated T84 cells and KFV119 or *rid::bla).*

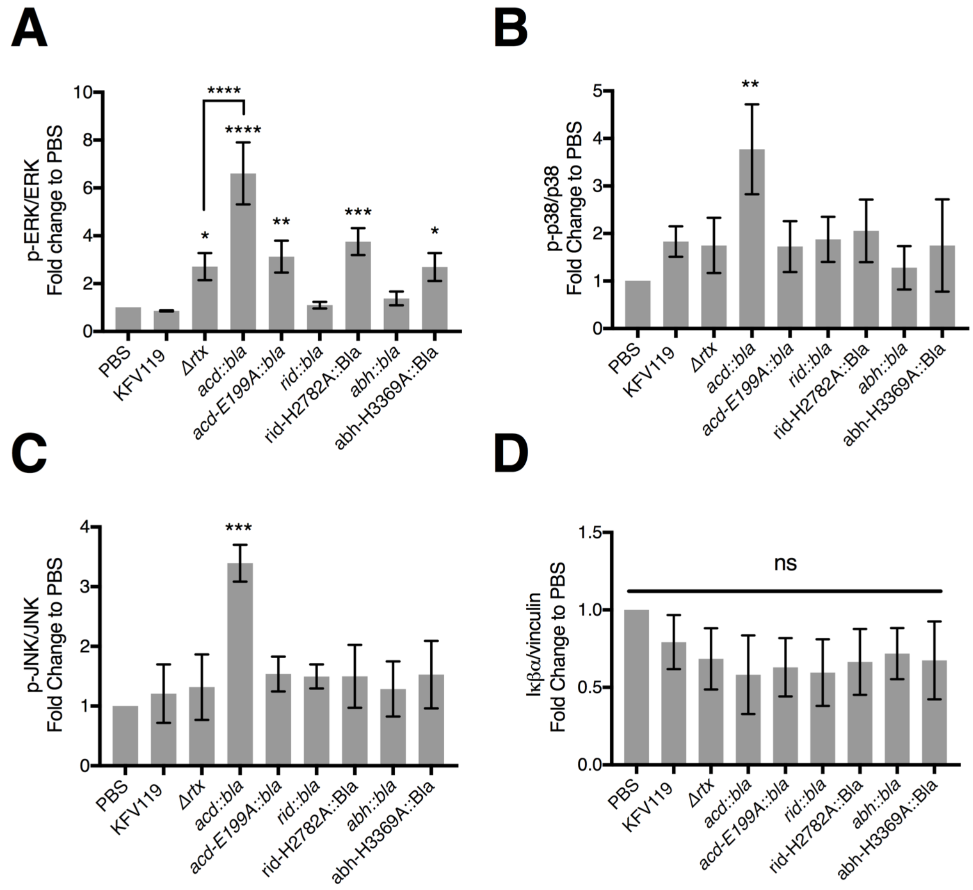


**Figure S2. ACD induced phosphorylation of ERK, p38, and JNK is suppressed when co-delievered with RID and ABH.**

(A–C) Densitometry analysis measuring fold change of phosphorylated ERK (p-ERK, A), phosphorylated p38 (p-p38, B), and phosphorylated JNK (p-JNK C) by the *V. cholerae* single effector gain-of-function strains as indicated. Data reported as mean ± s.d. $n = 3$ independent experiments (*p<0.05, ** p<0.01, ***p<0.001, One-way ANOVA with Tukey's multiple comparisons compared to the KFV119 WT strain or where indicated).

(D) Densitometry analysis measuring fold change of IκBα degradation. Data reported as mean ± s.d. $n = 3$ independent experiments.

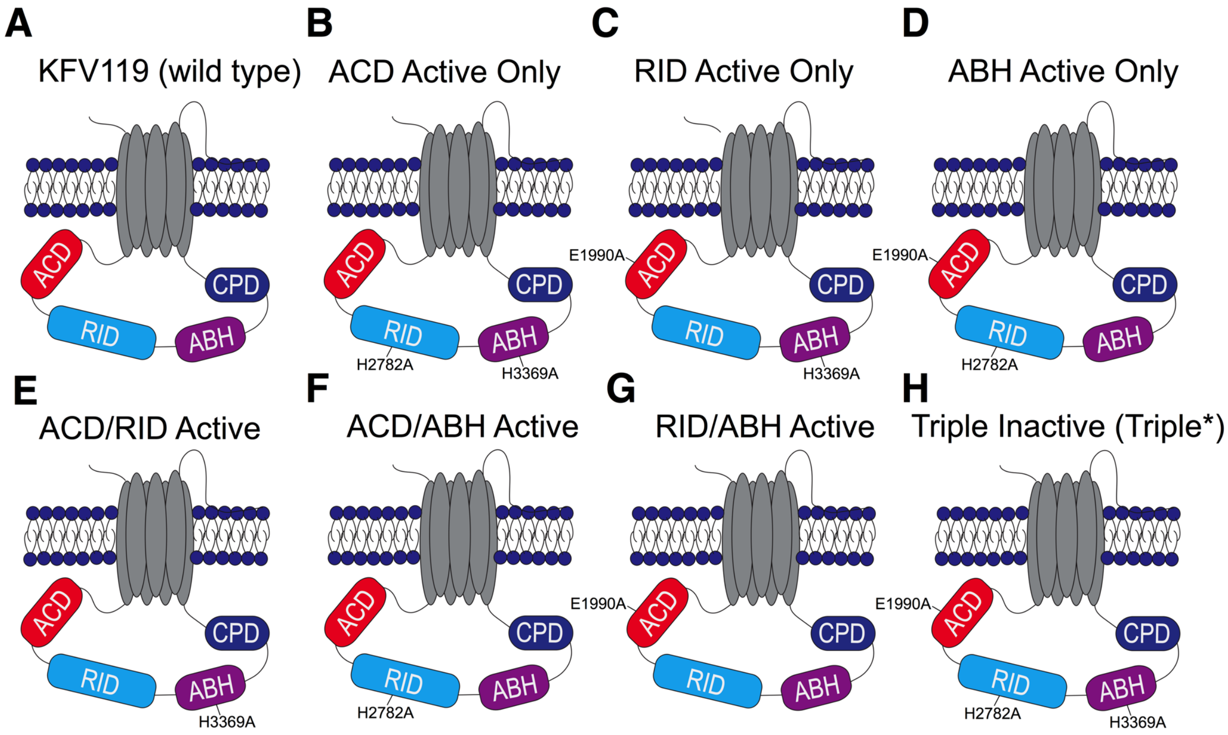


**Fig. S3. Schematic of single and double catalytically active MARTX$_{Vc}$ toxin effector strains and triple inactive MARTX$_{Vc}$ toxin effector strain.**

(A) Wild type toxin. (B) RID Active only strain with RID and ABH catalytically inactive. (C) RID Active only strain with ACD and ABH catalytically inactive. (D) ABH Active only strain with ACD and RID catalytically inactive. (E) ACD/RID active strain with only ABH catalytically inactive. (F) ACD/ABH Active strain with only RID catalytically inactive. (G) RID/ABH Active strain with only ACD catalytically inactive. (H) Triple inactive strain (Triple*) with all three effectors catalytically inactive.

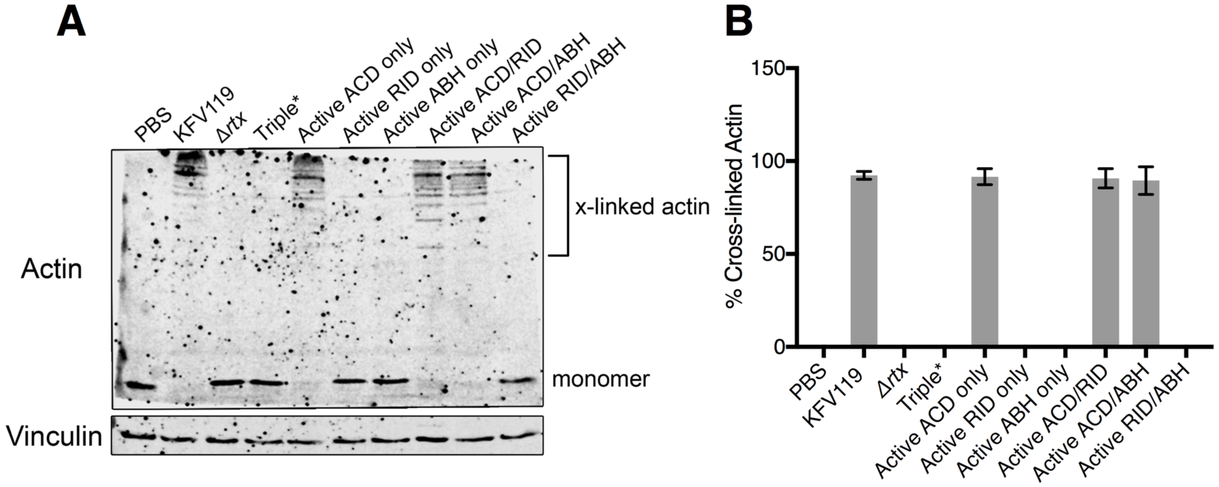


**Fig. S4. Actin laddering in T84 cells treated with V. *cholerae*.**

(A) Representative western blot of actin cross-linking from Hela cells inoculated with the constructed single and double catalytically active $MARTX_{Vc}$ toxin effector strains and triple inactive $MARTX_{Vc}$ toxin effector strain for 120 minutes and harvested in 2X SDS loading buffer.

(B) Quantification of actin cross-linking from western blot band density of actin cross-linking from Hela cells inoculated with the indicated strains normalized to KFV119. Data reported as mean ± s.d. $n$ = 3 independent experiments.

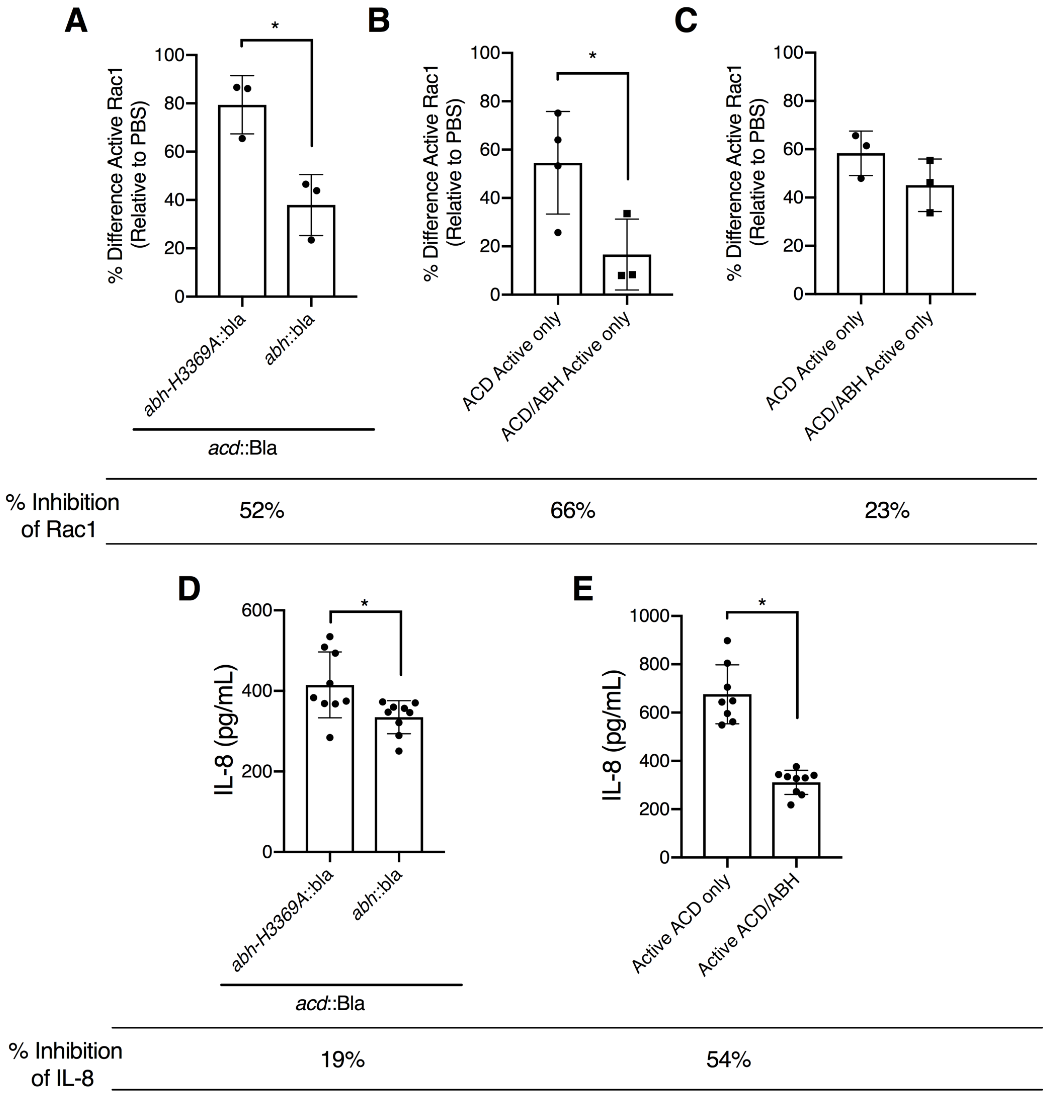


**Fig. S5. ABH mediated inhibition of host response to ACD varies across independent experiments.**

(A) Percent difference in Rac1 activation measured from T84 cells co-inoculated with the indicated Bla strains. Data from *n* = 3 samples reported as mean ± s.d. (*p<0.05, student's t-test).

(B-C) Percent difference in Rac1 activation measured from T84 cells inoculated with the indicated strains from two different independent sets of experiments (each set pooled data from *n* = 3 experiments). Data reported as mean ± s.d. (*p<0.05, Student's *t*-test).

(D-E) Comparison of ABH mediated suppression of ACD induced IL-8 secretion from T84 cells co-inoculated with *abh::bla* and *acd::bla* (D, data from Fig. 2G) or the ACD Active only and ACD/ABH

Active strains (E, data from Fig. 4B). Data pooled from $n = 3$ experiments reported as mean ± s.d. (*$p<0.05$, Student's *t*-test).

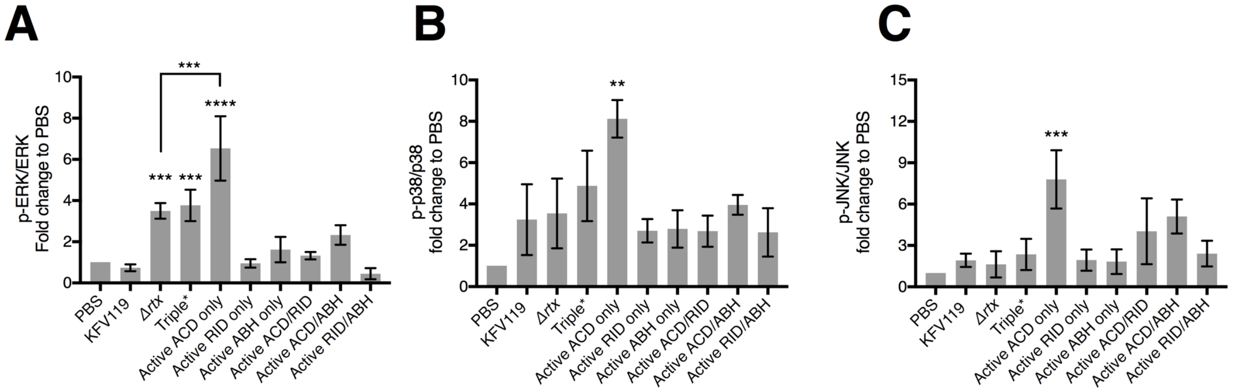


**Fig. S6. Both RID and ABH suppress ACD induced phosphorylation of ERK, p38, and JNK MAPK pathways.**

(A–C) Densitometry analysis measuring fold change of phosphorylated ERK (p-ERK, A), phosphorylated p38 (p-p38, B), and phosphorylated JNK (pJNK, C) by the *V. cholerae* single and double catalytically active $\text{MARTX}_{Vc}$ toxin effector strains and triple inactive $\text{MARTX}_{Vc}$ toxin effector strain as indicated. (*p<0.05, ** p<0.01, ***p<0.001, One-way ANOVA with Tukey's multiple comparisons compared to KFV119). *n* = 3 independent experiments.

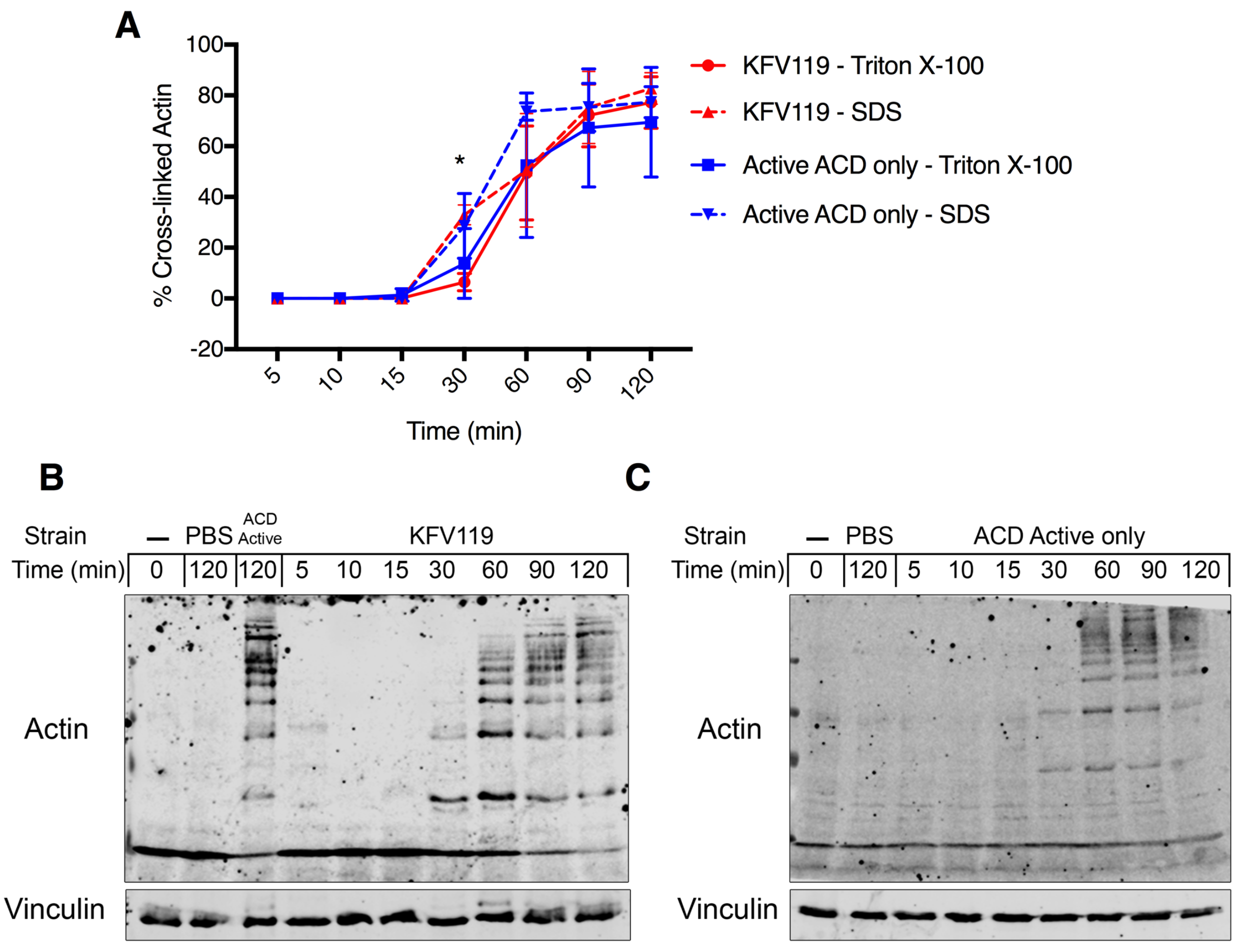


**Fig. S7. Actin crosslinking in cells harvested in Triton X-100 lysis buffer.**

(A) Quantification of actin cross-linking from T84 cells treated with KFV119 or the ACD Active Only strain with washed cells being lysed and collected in Triton X-100 lysis buffer followed by centrifugation and protein quantification or by direct resuspension and boiling in 2x SDS loading buffer. n = 3 independent experiments. Data from *n* = 3 reported as mean ± s.d.

(B) Western blot analysis of actin cross-linking from T84 cells inoculated with KFV119 and samples collected in SDS loading buffer. Blots representative of *n* = 3 independent experiments.

(C) Western blot of actin cross-linking form T84 cells inoculated with the ACD Active only strain and samples collected in SDS loading buffer. Blots representative of *n* = 3 independent experiments.

Note that Triton X-100 lysis underestimates actin cross-linking at early time points further supporting that extensive cross-linking activate the inflammatory response. Specifically, there is a statistically significant increase in actin cross-linkingobserved when KFV119 treated cells were lysed using SDS loading buffer compared to Triton X-100 at 30 minutes (*$p<0.05$, Two-way ANOVA with multiple comparisons). However, no significant difference was observed at any time point between the two methods when cells were treated with the ACD Active only strain. No significant difference was observed comparing samples treated with either KFV119 and the ACD Active only strain when using either method.

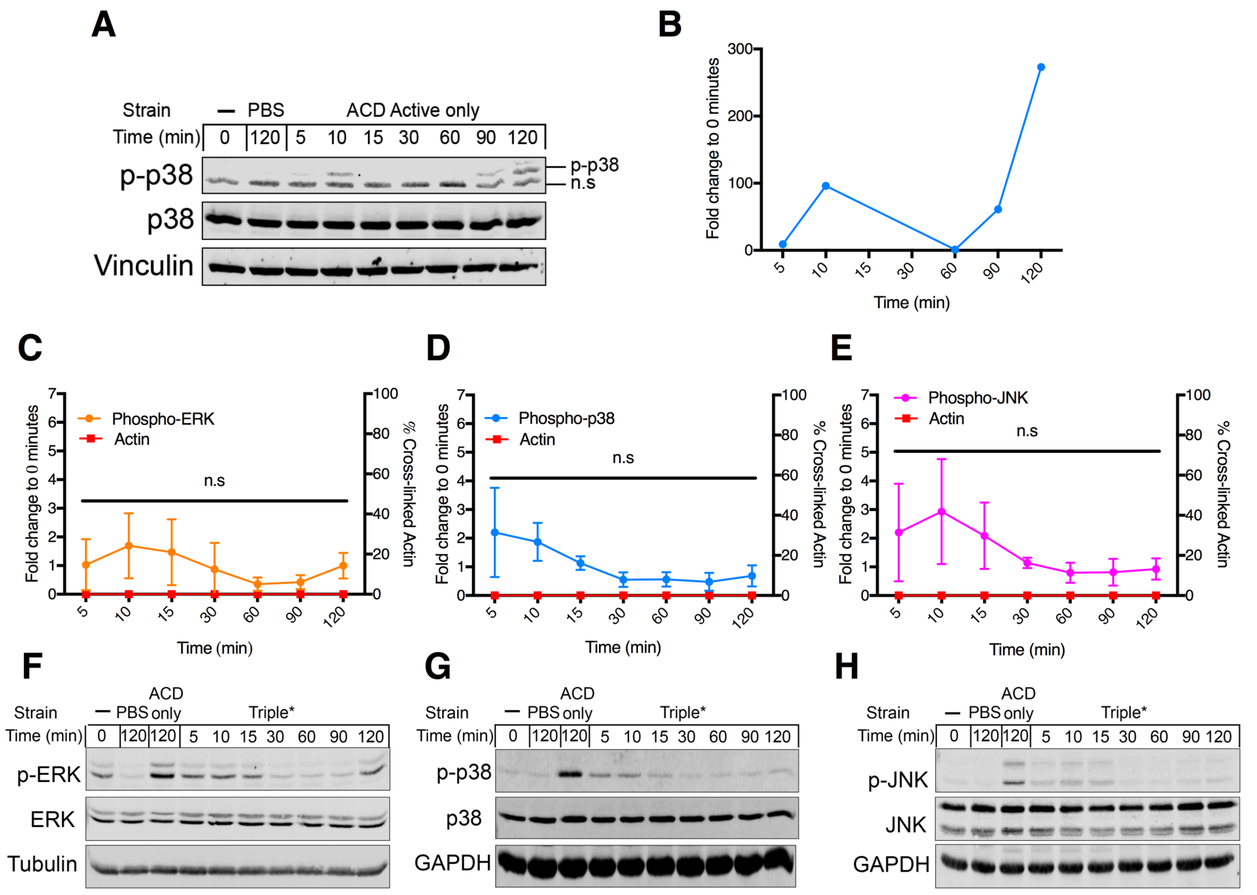


**Fig. S8. MAPK activation Triple* MARTX$_{Vc}$ toxin effector strain.**

(A) Western blot analysis of phosphorylated p-38 by the ACD only strain 273-fold times higher at 120 min compared to the 0 min control. Non-specific bands are indicated as n.s.

(B) Quantification of phosphorylated p-38 from western blot presented in panel A.

(C-E) Fold change of phosphorylated ERK (p-ERK, C), phosphorylated p38 (p-p38, D), and phosphorylated JNK (p-JNK, D) compared to 0 min control quantified from western blot band densities from T84 cells inoculated with the Triple* *V. cholerae* strain plotted with quantification of percent actin cross-linked over the two hour incubation. Data reported from $n$ = 3 experiments as mean ± s.d.

(F-H) Representative Western blot analysis ($n$=3 experiments) of phosphorylated ERK (p-ERK, F), phosphorylated p38 (p-p38, G), and phosphorylated JNK (p-JNK, H) activation during 120 min bacterial challenge.

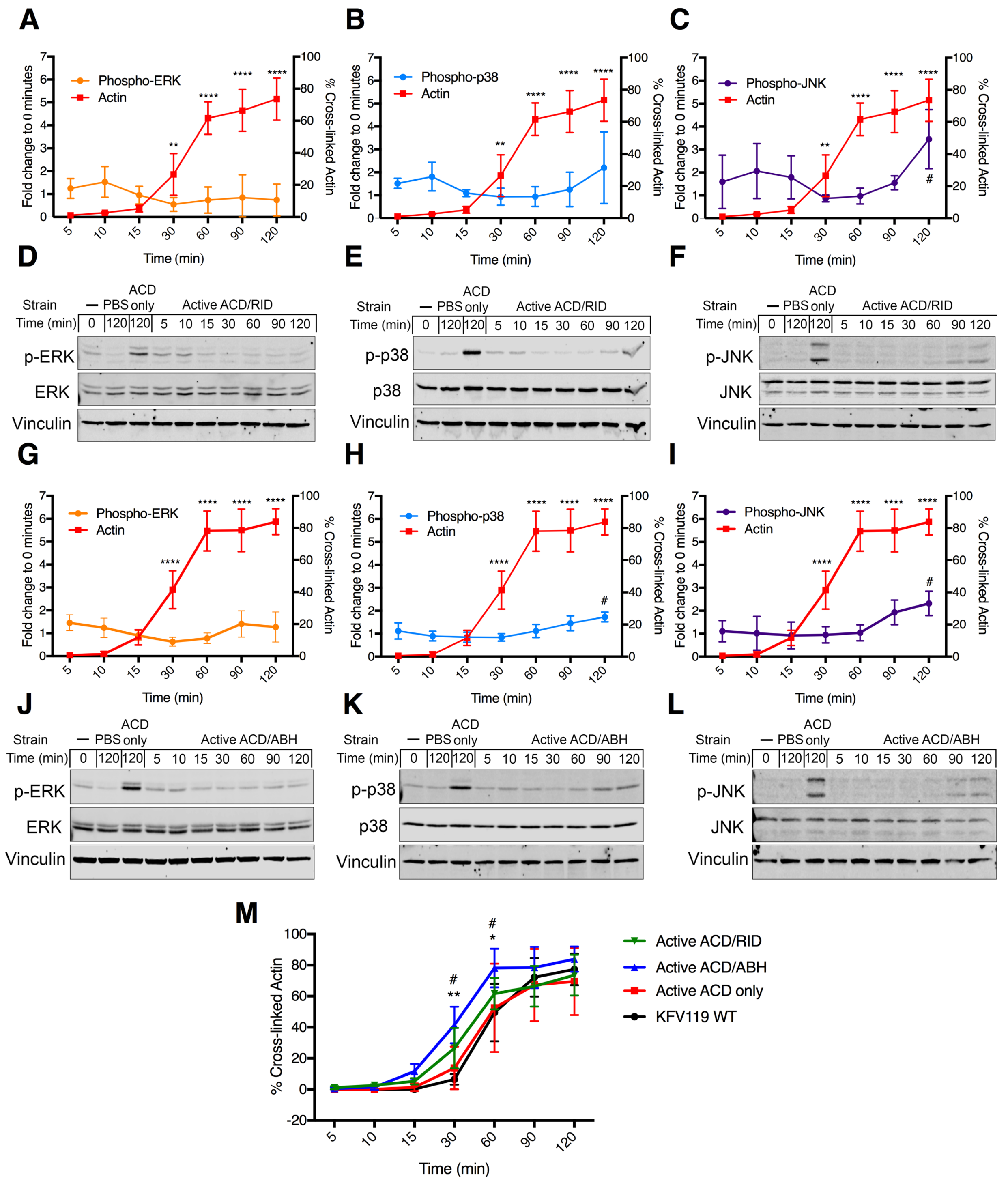


**Fig. S9. RID and ABH block MAPK signaling without modulating ACD activity.**

(A-C) Fold change of phosphorylated ERK (p-ERK, A) phosphorylated p38 (p-p38, B), and phosphorylated JNK (p-JNK, C) compared to 0 min control quantified from western blot band densities from T84 cells inoculated with the ACD/RID active *V. cholerae* strain plotted with quantification of percent actin cross-linked over the two hour incubation. Data reported from *n* = 4 (A), or 3 (B-C) experiments reported as mean ± s.d. (**p<0.01, ****p<0.0001, One-way ANOVA with Tukey's multiple comparisons between indicated timepoints and 0% actin cross-linking from 0 min control. # p<0.05, One-way ANOVA with multiple comparisons between phospho-JNK at indicated timepoint and a 0 min control with a normalized fold change of 1).

(D-F) Western blot analysis of phosphorylated ERK (p-ERK, D) phosphorylated p38 (p-p38, E), and phosphorylated JNK (p-JNK, F) activation during 120 min bacterial challenge. Blots representative of *n* = 4 (D), or 3 (E-F) experiments.

(G-I) Fold change of phosphorylated ERK (p-ERK, G) phosphorylated p38 (p-p38, H), and phosphorylated JNK (p-JNK, I) compared to 0 min control quantified from western blot band densities from T84 cells inoculated with the ACD/ABH active *V. cholerae* strain plotted with quantification of percent actin cross-linked over the 120 min incubation. Data from *n* = 3 (G-I) experiments reported as mean ± s.d. (****p<0.0001, One-way ANOVA with Tukey's multiple comparisons between indicated timepoints and 0% actin cross-linking from 0 min control. # p<0.05, One-way ANOVA with multiple comparisons between phosphorylated p38/JNK at indicated timepoints and a 0 min control with a normalized fold change of 1).

(J-L) Western blot analysis of phosphorylated ERK (p-ERK, J) phosphorylated p38 (p-p38, K), and phosphorylated JNK (p-JNK, L) activation during 120 min bacterial challenge. Blots representative of *n* = 3 (J-L) experiments.

(M) Comparison of percent actin cross-linking during 120 min bacterial challenge of ACD positive *V. cholerae* strains. Data quantified from *n* = 4 (Active ACD/RID) or 3 (all other strains) independent western blots and reported as mean ± s.d. (* p<0.05, ** p<0.01 Two-way ANOVA with multiple comparisons between KFV119 and Active ACD/ABH, #p<0.05, Two-way ANOVA with multiple comparisons between Active ACD only and Active ACD/ABH).

**Table S1. MARTX effector-dependent transcriptional responses in *V. cholerae*-infected T84 cells.**

Genes differentially regulated in T84 cells by the complete MARTX toxin, Δ*rtx*, or the single gain of function MARTX strains. This table is provided as an .xlsx file.

**Table S2. ACD-dependent transcriptional responses in in *V. cholerae*-infected T84 cells.**

Genes differentially regulated in T84 cells by the N16961 acd::bla strain compared to PBS-treated cells with a -1<= 0 <= 1 Log2 fold cut off for biological relevance. This table is provided as an .xlsx file.

**Table S3. Bacterial strains and plasmids used in study.**

| Designation | Relevant Description | Source |
|---|---|---|
| E. coli | | |
| BL21(DE3)(pMagic) | Protein overexpression, $Km^R$ | A. Joachimiak (Argonne National Laboratory) |
| DH5αλ*pir* | Plasmid cloning | Lab stock |
| SM10λ*pir* | Conjugation to *V. cholerae,* $Sm^R$ | Lab stock |
| V. cholerae | | |
| N16961Sm | El Tor O1, Wild-type, $Sm^R$ | (*63*) |
| KFV43 | N16961*ΔhapA* | (*64*) |
| VOV21 | N16961*ΔhlyA* | (*12*) |
| CW123 | N16961*ΔrtxA* | (*17*) |
| KFV119 | N16961*ΔhlyAΔhapA* | (*2*) |
| JD23 | KFV119*ΔrtxABCD* | (*2*) |
| JD1 | KFV119 *rtxA::bla* | (*2*) |
| JD20 | KFV119 *acd::bla* | (*2*) |
| JD19 | KFV119 *rid::Bla* | (*2*) |
| JD21 | KFV119 *rid-H2782A::bla* | (*2*) |
| JD2 | KFV119 *abh::bla* | (*2*) |
| JD15 | KFV119 *abh-H3369A::bla* | (*2*) |
| 2010EL-1786 | Atypical El Tor O1, Wild-type, $Sm^R$ | ATCC #BAA-2163 |
| JD16 | 2010EL-1786ΩpJD22 (2010EL-1786 *rtxA*+) | (*15*) |
| PJWV1 | KFV119 *rtxA-E1990A* (RID/ABH Active) | This Study |
| PJWV2 | KFV119 *rtxA-H2782A* (ACD/ABH Active) | This Study |
| PJWV3 | KFV119 *rtxA-H3369A* (ACD/RID Active) | This Study |
| PJWV4 | KFV119 *rtxA-E1990A-H2782A* (ABH Active Only) | This Study |
| PJWV5 | KFV119 *rtxA-E1990A-H3369A* (RID Active Only) | This Study |
| PJWV6 | KFV119 *rtxA-H2782A-H3369A (ACD Active Only)* | This Study |
| PJWV7 | KFV119 *rtxA-E1990A-H2782A-H3369A* (Triple*) | This Study |
| Plasmids | | |
| pTCO24 | Overexpression of $LF_N$-ACD, $Amp^R$ | (*36*) |
| pKS119 | Overexpression of $LF_N$-RID, $Amp^R$ | (*21*) |
| pSA129 | *sacB* counterselection cloning vector pWM91with *rtxA* with fragment of H2782A codon change, $Amp^R$ | (*55*) |
| pDS132 | *sacB* counterselection cloning vector, oriR6K, oriT, $Chl^R$ | (*65*) |
| pPJW4 | pDS132 with fragment of *rtxA* with E1990A codon change, $Chl^R$ | This Study |
| pPJW5 | pDS132 with fragment of *rtxA* with H3369A codon change, $Chl^R$ | This Study |

**Table S4. Sequences of gBlocks used  the *V. cholerae* genome.**

| Mutation | Sequence (5'-3') (codon change introduced in *V. cholerae* genome highlighted) |
|---|---|
| ACD E1990A | CCCAAGCTTCTTCTAGAGGTACCGCATGCCAAGCACAAGCCGATGCTCA<br>AGGTGCTAAACAAAACGAAGGTGATCGTCCTGATCGTCAAGGCGTGACT<br>GGTAGTGGCCTTTCGGGTAATGCTCATAGTGTGGAAGGCGCTGGCGAAA<br>CAGACAGTCATGTCAACACCGACAGCCAAACCAACGCCGATGGCCGATT<br>CAGTGAAGGTTTAACCGAACAAGAGCAAGAAGCGCTAGAAGGTGCGAC<br>CAACGCAGTGAACCGTTTGCAAATTAACGCAGGTATTCGAGCGAAAAAC<br>AGCGTTAGCAGTATGACTTCTATGTTCTCTGAAACAAATAGCAAGAGCAT<br>TGTTGTTCCTACCAAAGTCTCGCCTGAACCAGAGCGCCAAGAAGTGACTC<br>GTAGAGACGTCCGTATCTCAGGGGTGAACCTCGAAAGTCTAAGTGCGGT<br>ACAGGGAAGTCAACCAACGGGTCAACTGGCTTCGAAAAGTGTCCCCGGA<br>TTTAAAAGCCATTTCGCATCGACATCAATTGGTATA**GCA**AATGAGTTATC<br>CGGTCTGGTGGTGGTTTTACCGAAAAACTCAGCGCAGACTTTTGGCTATG<br>TGCATGATTCACAAGGTAACCCATTGTTCATGCTAACCAAGGATATGAAT<br>CAAGGTGGTTATAGCAACCCAGTGGGTATCAATGATATTCAAGGGGTGA<br>ACAACTGGCAGACGCATACGATTGAACTGGTTACATATCCTAGTGAAAT<br>CAGTGATACAGCAGCGGTTGAAAGTCGTAAAGAGGCAATGCTATGGCTT<br>GCGAAAGAGTTTACCGATCATATCAATCAGTCTAACCACCAAAGCTTACC<br>TCATTTAGTGAGTGATGACGGTCGTTTCACTCTGGTTATATCGAACTCTA<br>AGCATCTTATTGCGGCGGGTAACGGAACCTCTATTGATGCACAAGGCAA<br>GACCATAGGAATGACCCCTAGTGGCCAACAAGCAACAATGGCGATCAGT<br>GCGAAAGAATTTGGTACAAGCTCGTCGCCGGAAGTCAGACTGCTGCATG<br>CGATATCGAGCTCTCCCGGG |
| ABH H3369A | CCCAAGCTTCTTCTAGAGGTACCGCATGCAATCACTACCAGAAGCAAGG<br>TATCGATATGCTCGCAGTCAACCTGCGTGGCTATGGTGAAAGCGACGGT<br>GGACCAAGCGAAAAAGGCTTGTACCAAGATGCTCGCACCATGTTCAACT<br>ACCTAGTGAATGATAAGGGTATTGACCCAAGCAACATCATCATTCACGG<br>CTACTCAATGGGCGGTCCAATTGCCGCAGATTTAGCACGTTATGCCGCGC<br>AAAACGGCCAAGCGGTGTCTGGCTTATTGCTTGACCGTCCTATGCCAAGC<br>ATGACCAAAGCAATCACCGCTCACGAAGTGGCGAATCCAGCGGGCATTG<br>TGGGGGCTATCGCGAAAGCGGTTAACGGCCAGTTCTCTGTAGAGAAAAA<br>TCTCGAAGGTTTGCCAAAAGAGACATCCATTCTGCTGTTGACCGATAACG<br>AAGGTTTGGGTAACGAAGGTGAGAAACTTCGTACCAAACTCACTGCCTC<br>TGGTTACAACGTCACTGGCGAGCAGACATTCTATGGT**GCT**GAAGCAAGC<br>AACCGTTTGATGAGTCAATATGCGGATCAAATTGTCTCCGGTTTGTCCAG<br>CAGTGCAAGTGTAGATGAAGACCTAGATCAACAAGGGTTGGATACCACA<br>TCAACCAAGGATCAAGGTATCTCAAATAAGAATGATCATCTGCAGGTGG<br>TGGATAGTAAAGAAGCATTAGCGGATGGAAAAATACTCCATAATCAAAA<br>TGTTAATAGCTGGGGCCCGATTACGGTTACACCAACGACAGATGGTGGT<br>GAAACCCGCTTCGACGGTCAAATCATCGTTCAAATGGAAAACGACCCGG<br>TAGTAGCAAAAGCGGCAGCCAATTTAGCAGGTAAACATGCTGAAAGCAG<br>TGTGGTGGTGCAGCTCGATTCAGACGGCAACTATCGCGTGGTGTATGGCG<br>ATCCGTCAAAACTGGATGGAAAGCTACGTTGGCAGTTGGTGGGGCATGG<br>TCGCGACCACTCAGAAACTAACAATACTCGCTTAAGTGGTTACAGTGCCG<br>CATGCGATATCGAGCTCTCCCGGG |

**Table S5. Primers used in study.**

| Primer | Sequence | Function |
|---|---|---|
| IL-8_5' | AGCACTCCTTGGCAAAACTG | CXCL8 qPCR |
| IL-8_3' | CAAGAGCCAGGAAGAAACCA | CXCL8 qPCR |
| qTNF_5' | GCCAGAGGGCTGATTAGAGA | TNF qPCR |
| qTNF_3' | TCAGCCTCTTCTCCTTCCTG | TNF qPCR |
| qJUN_5' | GTCCTTCTTCTCTTGCGTGG | JUN qPCR |
| qJUN_3' | GGAGACAAGTGGCAGAGTCC | JUN qPCR |
| qFOS_5' | GGGGCAAGGTGGAACAGTTAT | FOS qPCR |
| qFOS_3' | CCGCTTGGAGTGTATCAGTCA | FOS qPCR |
| qCXCL3_5' | CGCCCAAACCGAAGTCATAG | CXCL3 qPCR |
| qCXCL3_3' | GCTCCCCTTGTTCAGTATCTTTT | CXCL3 qPCR |
| qEGR1_5' | AAAGCGGCCAGTATAGGTGA | EGR1 qPCR |
| qEGR1_3' | AGCCCTACGAGCACCTGAC | EGR1 qPCR |
| GAPDH_2_5' FWD | TTGAGGTCAATGAAGGGGTC | GAPDH qPCR |
| GAPDH_2_3' REV | GAAGGTGAAGGTCGGAGTCA | GAPDH qPCR |
| ACDVc_cat_FWD | GCACAAGCCGATGCTCAAGGTGCTAAACAAAAC | *acd* E1990A PCR |
| ACDVc_cat_REV | GTCTGACTTCCGGCGACGAGCTTGTACCAAATT | *acd* E1990A PCR |
| RIDVc_cat_FWD | GGCAAAGGTAATCTTGCCAATATCGATCTGCTAGG | *rid* H2782A PCR |
| RIDVc_cat_REV | CTCGATAAAGCGTTTCAGCTTAATGTCGCCATC | *rid* H2782A PCR |
| ABHVc_cat_FWD | CACTACCAGAAGCAAGGTATCGATATGTCGCAG | *abh* H3369A PCR |
| ABHVc_cat_REV | CCACTTAAGCGAGTATTGTTAGTTTCTGAG | *abh* H3369A PCR |
